# GreedyMini: Generating low-density DNA minimizers

**DOI:** 10.1101/2024.10.28.620726

**Authors:** Shay Golan, Ido Tziony, Matan Kraus, Yaron Orenstein, Arseny Shur

## Abstract

Minimizers is the most popular *k*-mer selection scheme in algorithms and data structures analyzing high-throughput sequencing (HTS) data. In a minimizers scheme, the smallest *k*-mer by some predefined order is selected as the representative of a sequence window containing *w* consecutive *k*-mers, which results in overlapping windows often selecting the same *k*-mer. Minimizers that achieve the lowest frequency of selected *k*-mers over a random DNA sequence, termed the expected density, are desired for improved performance of HTS analyses. Yet, no method to date exists to generate minimizers that achieve minimum expected density. Moreover, for *k* and *w* values used by common HTS algorithms and data structures there is a gap between the densities achieved by existing selection schemes and a recent theoretical lower bound. Here, we present `GreedyMini`, a toolkit of methods to generate minimizers with low expected or particular density, to improve minimizers, to extend minimizers to larger alphabets, *k*, and *w*, and to measure the expected density of a given minimizer efficiently. We demonstrate over various combinations of *k* and *w* values, including those of popular HTS methods, that `GreedyMini` can generate DNA minimizers that achieve expected densities very close to the lower bound, and both expected and particular densities much lower compared to existing selection schemes. Additionally, we show that the *k*-mer rank-retrieval time by `GreedyMini` is comparable to that of common *k*-mer hash functions. We expect `GreedyMini` to improve the performance of many HTS algorithms and data structures and advance the research of *k*-mer selection schemes.

## 1. Introduction

Minimizers are used in many bioinformatic applications, including sequence alignment, genome assembly, and data compression [1, 2, 3]. Minimizers are schemes that select representative sets of *k*-mers from sequences so that among every *w* consecutive *k*-mers the smallest *k*-mer according to some predefined order is selected. By efficiently identifying and indexing these *k*-mers, the computational complexity and memory requirements associated with processing large genomic datasets can be significantly reduced.

Minimizers are commonly evaluated by their expected or particular density. The expected density is the expected frequency of selected *k*-mers within a random DNA sequence, while the particular density is the frequency of selected *k*-mers in a specific DNA sequence [2]. Low density indicates a sparse and efficient selection, which enhances the performance of high-throughput sequencing (HTS) algorithms and data structures by reducing their runtime and memory usage. Thus, minimizers with low density are desired.

Several methods were developed to generate orders leading to low-density minimizers. Traditional lexicographical order © The Author 2022. Published by Oxford University Press. All rights reserved. For permissions, please often results in high density [4, 5]. Universal hitting sets (UHSs) were introduced to design orders for low-density minimizers. But, expensive heuristics are required to generate UHS-based orders limiting them to *k* ≤ 15 [6, 7, 8]. Decycling-based minimizers can be applied to any *k* value and achieve the lowest expected density compared to existing selection schemes to date [9]. Frequency-based orders can be used as a simple technique to generate minimizers with low particular density [10]. More recently, DeepMinimizer was developed to utilize machine learning to distribute selected *k*-mers more evenly, directly targeting low particular density [11]. To date, no method solves the problem of generating a minimum-expected-density minimizers scheme, and there is a gap between the theoretical lower bound and the densities achieved by existing selection schemes [12]. Some minimizers, such as Miniception [13] and syncmers [14], and other *k*-mer selection schemes, such as mod-minimizer [15] and strobemers [16], employ “internal” minimizers and can benefit from the use of low-density minimizers.

Here, we present `GreedyMini`, a toolkit of novel methods to generate minimizers with low expected or particular density, to locally improve a given minimizer, to extend minimizers to larger alphabets, *k*, and *w*, and to measure the expected density of a given minimizer efficiently. We prove density sguarantees for `GreedyMini`’s extensions to larger alphabets, and larger *k*. We combine these innovations into several pipelines to generate low-density DNA minimizers. We demonstrate over various combinations of *k* and *w* values, which are most practical for HTS analyses, that the pipelines generate minimizers that achieve lower density compared to existing selection schemes and very close to a recent theoretical lower bound [12] (e.g., within 1% for *w* = *k* − 1) and at comparable *k*-mer rank-retrieval time as common *k*-mer hash functions. We expect `GreedyMini` to improve the runtime and memory usage of HTS algorithms and data structures and to lead to further advancements in generating minimizers and *k*-mer selection schemes with low density.

## 2. Preliminaries

For integers *i* ≥ 0 and *j* ≥ *i* we denote [*i, j*) = {*i*,…, *j* − 1}. Let Σ = [0, *σ*) be an alphabet and *S* ∈ Σ^+^ be a non-empty string over Σ. *S*[*i*] denotes the letter at position *i* (the leftmost letter is *S*[0]), and *S*[*i, j*) refers to the substring covering the interval [*i, j*) of positions, so that *S* = *S*[0, |*S*|). Substrings of the form *S*[0, *j*) and *S*[*i*, |*S*|) are *prefixes* and *suffixes* of *S*, respectively. *Selection schemes* (defined below) operate by selecting substrings of small length *k*, called *k-mers*, in longer substrings called *windows*. We write *n-window* (and *n*-prefix, *n*-suffix, and *n*-string) to indicate length. Our algorithms assume the common unit-cost word-RAM model. If any integer function *g*(*σ, k, w*) appears as the time or space complexity of an algorithm, its values are assumed to fit into *O*(1) machine words; *σ* is treated as a constant. For a boolean expression *B*, [*B*] equals 1 if *B* is true and 0 otherwise.

A *local (selection) scheme* with parameters (*σ, k, w*) is a map *f*: Σ^*w*+*k*−1^ → [0, *w*). This map acts on strings over Σ, selecting one position in each (*w*+*k*−1)-window so that in a window *v* = *S*[*i, i* + *w* + *k* − 1) the position *i* + *f* (*v*) in *S* is selected.

The most popular metric to evaluate a local scheme by is the density of selected positions. Let *f* (*S*) denote the set of positions selected in a string *S* by a local scheme *f*. The *particular density of f on S* is 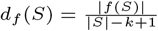. The *expected density of f* is the limit 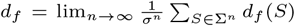. The *density factor* of *f* is a normalization of *d*_*f*_ by a factor of (*w*+1), i.e., *df*_*f*_ = (*w* + 1)*d*_*f*_.

Let *π* be a permutation (= a linear order) of the set of all *σ*-ary *k*-mers. We view *π* as a bijection *π*: [0, *σ*^*k*^) → Σ^*k*^ and say that a *k*-mer *u* has *rank i*, denoted by `rank` _*π*_(*u*) = *i*, if *π*(*i*) = *u*. The *minimizer* (*π, w*) is a local scheme that maps each (*w*+*k*−1)-window to the starting position of its minimum-rank *k*-mer with ties broken to the left.

A subset *H* ⊆ Σ^*k*^ is a *universal hitting set* (UHS) for *w* if every (*w*+*k*−1)-window contains at least one *k*-mer from *H*. Let *π* be a linear order on Σ^*k*^ and let *i* be such that *H* = {*u* | `rank` _*π*_(*u*) *< i*} is a UHS for *w*. Every permutation *π*^*′*^ such that *π*^*′*^(*j*) = *π*(*j*) for all *j* ∈ [0, *i*) generates the same minimizer: (*π*^*′*^, *w*) = (*π, w*). Accordingly, we define a *UHS order on* Σ^*k*^ (for *w*) to be any bijection *ρ*: [0, |*H*|) → *H*, where *H* ⊆ Σ^*k*^ is a UHS for *w*. Any completion of *ρ* to a permutation *π* of Σ^*k*^ defines the same minimizer, which we denote by (*ρ, w*). We define `rank` _*ρ*_(*u*) as above if *u* ∈ *H* and set `rank` _*ρ*_(*u*) = ∞ otherwise. Since Σ^*k*^ is a UHS for any *w*, a permutation of Σ^*k*^ is a special case of a UHS order.

For a given minimizer *f* = (*ρ, w*), a (*w*+*k*)-window *v* is *charged* if its minimum-rank *k*-mer is either its prefix or its *unique suffix* (i.e., the *k*-suffix of *v* having no other occurrence in *υ*); otherwise, *υ* is *free*. An important observation is that every string *S* contains exactly |*f* (*S*)| − 1 (not necessarily distinct) charged (*w*+*k*)-windows (Lemma 6 in [2]). Since all possible *n*-strings have, in total, the same number of occurrences of each (*w*+*k*)-window, the expected density *d*_*f*_ of a minimizer equals the fraction of charged windows in Σ^*w*+*k*^ [5, 13].

## 3. Methods

### 3.1. Generating minimizer orders greedily

We present a novel randomized algorithm `GreedyE` (Algorithm 1, `E` stands for expected) to generate low-density minimizers. Given parameters (*σ, k, w*), `GreedyE` generates a UHS order *ρ* on Σ^*k*^ for *w*, assigning ranks to *k*-mers one by one, starting from rank 0 and stopping when the *k*-mers with assigned ranks constitute a UHS. `GreedyE` maintains a score function for all unranked *k*-mers. At each iteration, `GreedyE` creates a pool of low-scored *k*-mers, chooses a random *k*-mer from it, assigns the next rank to the chosen *k*-mer, and updates the scores of all unranked *k*-mers.

During an iteration, a (*w*+*k*)-window is *live* if all its *k*-mers are unranked. For an unranked *k*-mer *u*, let *X*_*u*_ be the set of all live windows containing *u*, and let *Y*_*u*_ ⊆ *X*_*u*_ consist of live windows containing *u* as a prefix or as a unique suffix. Then, `score` (*u*) = |*Y*_*u*_|*/*|*X*_*u*_|. The intuition behind selecting a low-scored *k*-mer is the following: choosing *u* to have the next rank clarifies the status (free or charged) of the windows from *X*_*u*_ and only of them, and those from *Y*_*u*_ are charged. Thus, `score` (*u*) locally approximates the fraction of charged windows among all windows, which is the fraction we want to minimize.

#### Algorithm 1: GreedyE: Generating a low-density UHS order

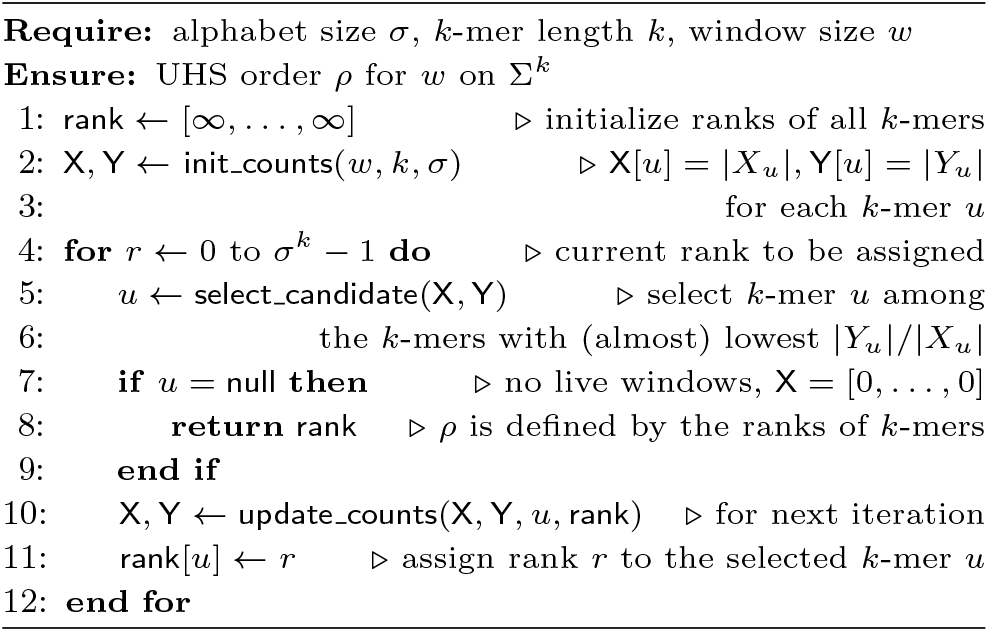

In our implementation, we define low-scored *k*-mers as follows. For each run of `GreedyE` we sample a parameter *α* from a user-defined range: *α* ∼ LogUniform(*α*_*min*_, *α*_*max*_). At each iteration of `GreedyE`, an unranked *k*-mer *u* is considered low-score if its *inverted score* 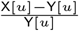 is at least *αI*, where *I* is the maximum inverted score at this iteration. With *α*_*min*_ = *α*_*max*_ = 1, `GreedyE` selects a lowest-score *k*-mer at each iteration but still uses randomization when several *k*-mers are tied for the lowest score. Decreasing *α*_*min*_ and *α*_*max*_ expands the search space of low-density minimizers at no additional space or runtime costs, and thus has the potential to improve the best result over multiple runs.

The natural computational requirements for implementing `GreedyE` are: Ω(*σ*^*k*^) space to store the order; Ω(*σ*^*k*^) time per iteration to select a low-score *k*-mer; Ω(*wσ*^*k*+*w*^) time to scan each window once. Theorem 1 presents our implementation, which meets all these lower bounds up to additional *O*(*w*) space. We prove the theorem and give the pseudocode of non-trivial auxiliary functions in Supplementary Section S4.

#### Theorem 1

`GreedyE` *can be implemented to run in O*(*σ*^*k*^ + *w*) *space and O*(|*H*|(*σ*^*k*^ +*wσ*^*w*^)) *time, where H is the constructed UHS*.

#### Remark 1

*(i) The size of every UHS is between* 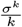 *and σ*^*k*^. *(ii)* `GreedyE` *can be implemented with the running time O*(|*H*|*wσ*^2*k*^), *which is linear in w but has heavier dependence on k. We found no practical use for this implementation, so we only briefly describe it in Supplementary Section S4*.

To generate minimizers of low *particular* density with respect to a specific input string *S*, we present an analog of `GreedyE`, called `GreedyP`. In this case, each window has a weight equal to the number of its occurrences in *S*. `GreedyP` computes and stores the windows occurring in *S* with their weights at an additional expense of *O*(|*S*|) time and *O*(min{|*S*|, *σ*^*w*+*k*^}) space, since the number of (*w* + *k*)-windows of positive weight is bounded by the observed windows in *S* and by *σ*^*w*+*k*^. Then, the only difference with `GreedyE` (Algorithm 1) is that the arrays `X` and `Y` contain *sums of weights* of windows rather than the numbers of windows. For the case where |*S*| ≪ *σ*^*w*+*k*^, `GreedyP` can be implemented more efficiently based on a linked list of windows with positive weight.

### 3.2. Improving minimizers by hill climbing

We apply a randomized version of the standard local search technique called *hill climbing* to improve a given minimizer (*ρ, w*), where *ρ* is a UHS order for *w*. At each iteration, the search algorithm samples a random rank *r* such that both *r* and *r*+1 are assigned in *ρ* and calls the function `Swap` (*ρ, w, r*), defined as follows. `Swap` (*ρ, w, r*) generates the UHS order *ρ*^*′*^from *ρ* by swapping the *k*-mers with ranks *r* and *r*+1 and calculates 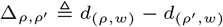. If 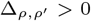, `Swap` replaces *ρ* with *ρ*^*′*^; if 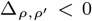, `Swap` does nothing; if 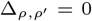, `Swap` replaces *ρ* with *ρ*^*′*^ with probability 0.5. The search stops when a user-defined time limit or number of iterations is reached.

Theorem 2 presents the complexity of our two efficient implementations of `Swap`. Our default implementation (a) uses recursive depth-first traversal (`SwapDFS`), while (b) is designed for the case of large *w* and based on dynamic programming (`SwapDP`). The proof of Theorem 2 and the pseudocodes of `SwapDFS` and `SwapDP` are presented in Supplementary Section S3.

#### Theorem 2

*Let ρ be a UHS order on* Σ^*k*^ *for w, and let the ranks r, r*+1 *be assigned in ρ. The function* `Swap` (*ρ, w, r*) *can be implemented in either (a) O*(*σ*^*k*^ + *σ*^*w*^) *time and O*(*w*) *additional space or (b) O*(*wσ*^*k*^) *time and O*(*σ*^*k*^) *additional space*.

### 3.3. Extending orders to larger *σ, k*, and *w*

A local scheme *f* with the parameters (*σ, w, k*) can be extended to a local scheme *f*^*′*^ with the parameters (*σ, w, k*^*′*^), where *k*^*′*^ *> k*, by a simple rule [17]: in every (*w*+*k*^*′*^−1)-window, *f*^*′*^ picks the position chosen by *f* in the (*w*+*k*−1)-prefix of this window. Trivially, 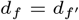. In a similar way, a local scheme *f*_*γ*_ with the parameters (*σγ, w, k*) can be built from *f*. Namely, the alphabet is partitioned into *σ* classes, and in every window *υ f*_*γ*_ picks the position chosen by *f* in the window 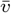, obtained from *υ* by replacing each letter with its class. Here as well,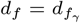.

Such extensions may suffice for practice. But, their main theoretical limitation is that *f*^*′*^ and *f*_*γ*_ do not inherit the property “to be a minimizer” from *f*. Namely, if *f* is a minimizer, then *f*_*γ*_ is never a minimizer, while the status of *f*^*′*^ depends on *f* and *w*. In particular, if *w > k*^*′*^, then *f*^*′*^ is not a minimizer for any *f*. See Supplementary Section S5 for a detailed discussion.

We upgrade these trivial extensions to obtain minimizer-to-minimizer extensions, and prove their density guarantees. We define *dual minimizers* as local schemes that act like minimizers but break ties to the right. For a minimizer *f*, we denote the dual minimizer using the same order as *f* by *f*^←^. Notice that *d*_*f*_ and 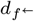 are close: only the windows containing their minimum-rank *k*-mer as a non-unique prefix or a non-unique suffix are charged differently (e.g., the average difference in expected density was less than 0.15% over 1000 random minimizers for *σ* = 2, *k* = 8, and *w* = 12). Given a UHS order *ρ* for *w* on Σ^*k*^, we define its *mirror* as the UHS order *ρ*^←^ such that 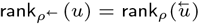 for all *u* ∈ Σ^*k*^, where 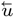 is *u* reversed (Figure 1). Lemma 1 connects mirrors to dual minimizers.

**Fig. 1.**
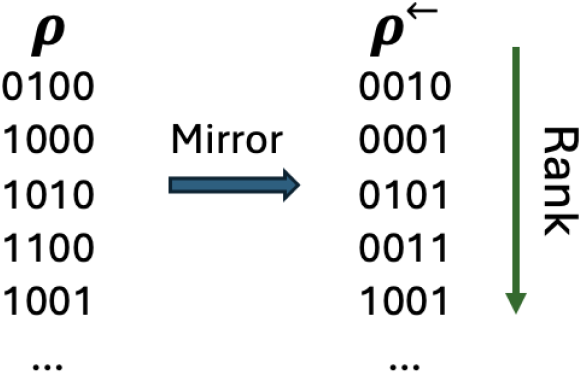
An example of a binary order *ρ* on 4-mers and its mirror *ρ*^←^.

#### Lemma 1

If *ρ* is a UHS order for *w*, then 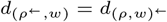.

*Proof* If a (*w*+*k*)-window *υ* is charged by the scheme (*ρ, w*)^←^, its *k*-mer *u* with the minimum *ρ*-rank is either a suffix or a unique prefix of *υ*. Hence, in the window 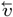 the *k*-mer 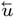 has the minimum (*ρ*^←^)-rank and is either a prefix or a unique suffix; respectively, 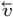 is charged by (*ρ*^←^, *w*). As all steps are reversible, we get a bijection between the sets of charged windows of the two schemes. □

We transform an arbitrary minimizer *f* with the parameters (*σ, k, w*) to a minimizer *f*^*′*^ with the parameters (*σγ, k, w*) for an integer *γ >* 1 as follows. Let Γ = [0, *γ*) and consider the alphabet 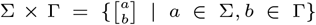 of size *σγ*. Every string 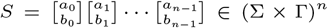 has two *projections S*_Σ_ = *a*_0_…*a*_*n*−1_ ∈ Σ^*n*^ and *S*_Γ_ = *b*_0_…*b*_*n*−1_ ∈ Γ^*n*^. Given a UHS order *ρ* for *w* on Σ^*k*^ and a linear order *τ* on Γ^*k*^, we define a UHS order *ρ × τ* for *w* on (Σ *×* Γ)^*k*^ by setting `rank` _*ρ×τ*_ (*u*) = *γ*^*k*^`rank` _*ρ*_(*u*_Σ_) + `rank` _*τ*_ (*u*_Γ_); in particular, `rank` _*ρ×τ*_ (*u*) = ∞ iff `rank` _*ρ*_(*u*_Σ_) = ∞. The following theorem presents a minimum density guarantee over the minimizers (*ρ×τ, w*) and (*ρ*^←^*×τ, w*). Its proof is provided in Supplementary Section S6.

#### Theorem 3

*Let ρ be a UHS order on* Σ^*k*^ *for w and τ be a linear order on* Γ^*k*^. *Then*, min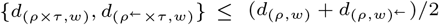.

We use a similar approach to obtain the same density guarantee for the extension of a minimizer with parameters (*σ, k, w*) to a minimizer with the parameters (*σ, k*+1, *w*). Given a UHS order *ρ* on Σ^*k*^ for *w*, we define its *extensions ρ*_1_ and *ρ*_2_ on Σ^*k*+1^ as follows. For *u* ∈ Σ^*k*^ and *a* ∈ Σ, we set 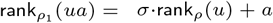 and 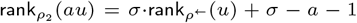. We prove the next theorem in Supplementary Section S7.

#### Theorem 4

*Let ρ be a UHS order on* Σ^*k*^ *for w, and let ρ*_1_ *and ρ*_2_ *be its extensions. Then*, min 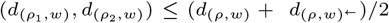.

### 3.4. Measuring the density of a given minimizer

A standard approach to measure the expected density of a given selection scheme is to generate a long (pseudo)random string and count sampled positions. However, the exact calculation of the expected density is preferred whenever it is feasible. As was pointed out in [5], the exact density can computed by processing a de Bruijn sequence of order *w* + *k* instead of a random string, as such a de Bruijn sequence contains each (*w* + *k*)-window exactly once. In this way, the expected density can be computed in *O*(*σ*^*w*+*k*^) time and *O*(*w* + *k*) space using a fast shift rule for de Bruijn sequences [18, 19].

We present two new algorithms, `DenDFS` and `DenDP`, that calculate the exact expected density of a given minimizer more efficiently. Their complexity is given by Theorem 5 (a) and (b), respectively. The algorithms and the proof of Theorem 5 are presented in Supplementary Section S2.

#### Theorem 5

*Let ρ is a UHS order on* Σ^*k*^ *for w and let H be its UHS. The expected density of the minimizer* (*ρ, w*) *can be computed in either (a) O*(|*H*|*σ*^*w*^) *time and O*(*w*) *space, or (b) O*(*w*|*H*|*σ*^*k*^) *time and O*(*σ*^*k*^) *space*.

#### Remark 2

*(i) Since the size of H is at most σ*^*k*^ *and can be as small as* 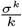, `DenDFS` *is faster than the algorithms that process a de Bruijn sequence. In our benchmarking (Section 4*.*1)*, *more than* 90% *of the binary UHS orders satisfy* 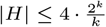. (*ii*) `DenDP` *avoids exponential dependence on w and thus computes the density for very large windows at a modest space expense*.

### 3.5. GreedyMini toolkit and pipelines

We combined the novel techniques from Sections 3.1–3.4 into a toolkit called `GreedyMini`. Its primary purpose is the generation of low-density DNA minimizers for given parameters *k* and *w*.

The main tools are `GreedyE` and `GreedyP` (Section 3.1), implemented for the *binary* alphabet. `GreedyE` takes the parameters *k, w*, a range (*α*_*min*_, *α*_*max*_), and returns a UHS order *ρ* on {0, 1}^*k*^ for *w* such that the minimizer (*ρ, w*) has low expected density. `GreedyP` has the same input parameters plus a sequence *S*, and outputs a UHS order *ρ* on {0, 1}^*k*^ such that the minimizer (*ρ, w*) has low particular density for *S*.

The tools `SwapDFS` and `SwapDP` (Section 3.2) take the parameter *w* and a UHS order *ρ* on {0, 1}^*k*^ for *w* as input and return an order *ρ*^*′*^ on {0, 1}^*k*^ such that 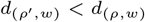, unless no such order *ρ*^*′*^ was found, and then *ρ* is returned. By default, `SwapDFS` is used, while `SwapDP` is intended to cover large values of *w*.

The tools `DenDFS` and `DenDP` (Section 3.4) compute the exact expected density of *binary* minimizers with `DenDFS` being the default method and `DenDP` covering large values of *w*.

Finally, the toolkit includes auxiliary functions described in Section 3.3. `Mirror` receives a binary UHS order *ρ* and returns the order *ρ*^←^; `Extend` -*k* receives a binary minimizer and extends it to a larger value of *k*; `Extend` -*σ* receives a binary minimizer and extends it to the DNA alphabet (taking *σ* = *γ* = 2 in the extension definition, with *τ* being the lexicographic order).

We implemented several pipelines as part of `GreedyMini`.

#### GM *-*expected: *Generating a DNA minimizer for* (*k, w*)

To construct a DNA minimizer with low *expected* density for given *k* and *w*, we proceed in three stages (Supplementary Figure S2A).

##### Generate

Given *k* and *w*, run multiple instances (by default 4096) of `GreedyE` (2, *k, w*); the default range (*α*_*min*_, *α*_*max*_) to sample the parameter *α* is (0.94, 0.9996). Report a lowest-density minimizer or its mirror, whichever has lower density.

##### Improve

For the minimizer reported by stage Generate, run an instance of `SwapDFS` on each available CPU core (64 on our machine) with the time limit set to the runtime of stage Generate (in our experiments, this resulted in *>*400 · 2^*k*^ `Swap` calls per core). Report a lowest-density minimizer.

##### Extend

Given the minimizer (*ρ, w*) from stage Improve, report the DNA minimizer `Extend` -*σ*(*ρ, w*) or `Extend` -*σ*(*ρ*^←^, *w*), whichever achieves lower density.

#### GM *-*particular: *Generating a DNA minimizer for* (*k, w*) *and sequence S*

To construct a DNA minimizer with low *particular* density for given *k, w*, and a sequence *S*, we perform stage Generate, replacing `GreedyE` by `GreedyP`. Then, we skip stage Improve and run stage Extend to obtain the result (Supplementary Figure S2B).

#### GM *-*improve: *Generating a DNA minimizer for large w*

If running `GM` -`expected` on the pair (*k, w*) is expected to take too much time due to large *w*, we start with a binary minimizer obtained for the pair (*k, w*^*′*^) for some *w*^*′*^ *< w*. We run an instance of `SwapDP` on each available CPU core within a manually set time limit (10 hours by default). Finally, we report the lowest-density minimizer over these runs. For the reported minimizer, we run stage Extend to obtain the final result (Supplementary Figure S2C). Additionally, `GM` -`improve` can be used to improve any binary minimizer via the use of `SwapDP`.

#### GM *-*k: *Generating a DNA minimizer with increased k*

If running `GM` -`expected` on the pair (*k, w*) resulted in a minimizer having higher density than the minimizer obtained for the pair (*k* − 1, *w*), we start with the binary minimizer obtained for (*k* − 1, *w*), apply `Extend` -*k* on it, and run stage Improve within a manually set time limit. Then, we run stage Extend to obtain the result. This pipeline is useful for the case where *k* is large enough and *w* is small enough, as running `Swap` takes just *O*(2^*w*^) time, while the runtime of an instance of `GreedyE` is *O*(2^2*k*^).

#### Trivial extensions

If running `SwapDP` and `SwapDFS` for the pair (*k, w*) takes too much time or is even infeasible, we use trivial extensions. For a trivial *k*-extension, we take a binary minimizer for some parameters *k*^*′*^ *< k* and *w*, apply `Extend` -*k*, and run stage Extend on it. For a trivial *w*-extension, we take a binary order *ρ* obtained for some parameters *k* and *w*^*′*^ *< w*, consider the minimizer (*ρ, w*), and run stage Extend on it.

## 4. Results

### 4.1. Benchmarking *k*-mer selection schemes

We used `GreedyMini` to generate low-density DNA minimizers for various *k, w* combinations and benchmarked the results against the minimizers constructed by the state-of-the-art methods: double decycling [9], miniception [13], and the recent open-closed syncmer [20], which mixes the ideas of the previous two. For these methods we used the implementations from the mod-minimizer GitHub repository [21] and measured their density using a random 4-ary 10^7^-string. We excluded from the comparison decycling [9], DOCKS [7], and PASHA [8] methods as they are clearly outperformed by double decycling [9].

To benchmark in the mode “fixed *w*, variable *k*”, we took *w* = 8 and *w* = 12. For *w* = 8, we ran `GM` -`expected` for all *k* ∈ [3, 18) and made a trivial *k*-extension up to *k* = 30. For *w* = 12, we ran `GM` -`expected` for all *k* ∈ [3, 17) and extended trivially up to *k* = 26. Similarly, to benchmark in the mode “fixed *k*, variable *w*”, we took *k* = 8 and *k* = 12. For *k* = 8, we ran `GM` -`expected` for all ∈ [3, 20), and made a trivial *w*-extension up to *w* = 100. For *k* = 12, we ran `GM` -`expected` for all ∈ [3, 17), and then ran `GM` -`improve` subsequently for *w* = 20, 25, 30, taking the previous result as the starting point. We then trivially extended the resulting minimizer up to *w* = 100.

In addition, we ran `GM` -`expected` over all (*k, w*) pairs with 3 ≤ *k, w* ≤ 15 and focused on the mode “*w* ≈ *k*”. There were a few cases where the density obtained by `GM` -`expected` for (*k, w*) was higher than the one obtained for (*k* − 1, *w*). In each such case, we ran `GM` -`k` to improve the result.

Finally, we addressed specific (*k, w*) pairs used in popular HTS methods: (9, 16) (KMC 3) and (15, 17) (Kraken) were processed by `GM` -`expected`; for (7, 22) and (7, 49) (both KMC 2) we made a trivial *w*-extension from *w* = 15; for (21, 11) (SSHash), (29, 11) (Giraffe), and (31, 5) (Kraken 2) we made a trivial *k*-extension from *k* = 15; for (19, 30) (GraphAligner) we made a trivial *k*-extension from *k* = 12.

The expected density of the resulting DNA minimizers was computed as follows: (i) *k* + *w* ≤ 26, except for trivial extensions: exact computation by `DenDFS` /`DenDP`; (ii) 27 ≤ *k* + *w* ≤ 30, except for trivial extensions: the binary density is computed exactly, and the number of bad/good (*k* + *w*)-windows (Supplementary Section S6) is estimated from a sample of 10^8^ windows over Γ; (iii) *k* + *w* ≥ 31, plus all trivial extensions: the binary density is computed exactly, then the upper bound of Theorem 3 is reported.

We also ran `GM` -`particular` to generate minimizers with low particular density for a DNA sequence of 10^6^ nucleotides from Chromosome X for *w* = 12 and all *k* ∈ [3, 17), and for *k* = 8 and all *w* ∈ [3, 20). We compared the results to the same set of benchmarked selection schemes; our attempts to compare to DeepMinimizer [11] and polar sets [22] failed, because their GitHub implementations raised errors and lacked documentation (Supplementary Section S9). We report the minimizer construction runtime and maximum memory usage of `GM` -`expected` and `GM` -`particular` in Supplementary Section S10.

### 4.2. Expected and particular density results

For fixed *w* (Figures 2A,B), `GM` -`expected` outperformed all other minimizers, losing to open-closed syncmer in a single point (12, 26). For *k* = 8 (Figure 2C), `GM` -`expected` outperforms all other minimizers by a large margin, even though it uses the trivial *w*-extension.. For *k* = 12 (Figure 2D), trivial *w*-extension of the `GM` -`expected` from *w* = 16 is not enough: its result is nearly tied with the result of double decycling for *w >* 20. However, the use of `GM` -`improve` results in a clear gap of 2.40% − 3.86%. We observed similar trends for particular density (Figures 2E,F), where the best performers were `GM` -`particular` and `GM` -`expected`, with `GM` -`particular` achieving clearer superiority for *w* + *k >* 24.

**Fig. 2.**
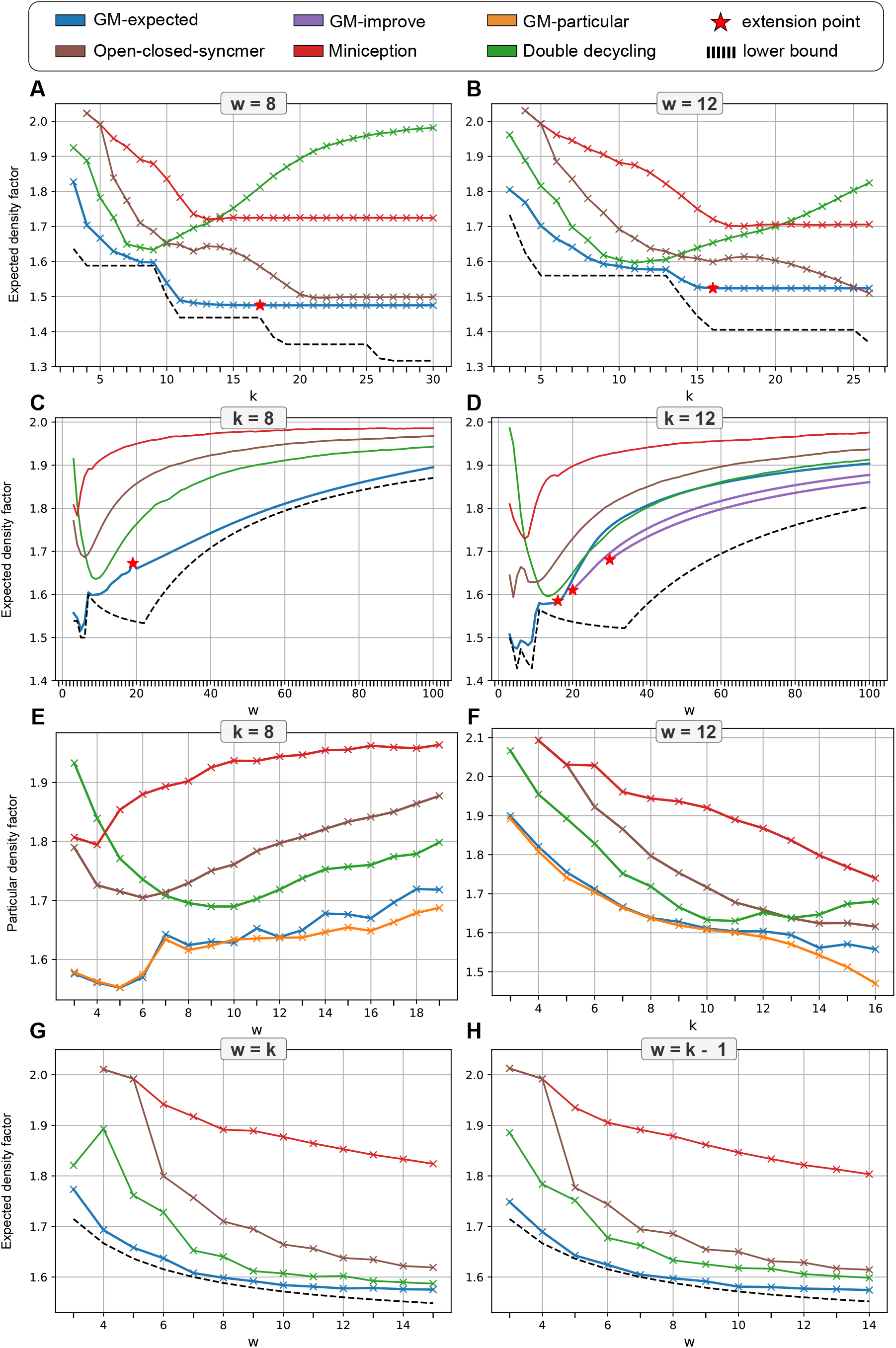
Performance of various DNA *k*-mer selection schemes over various (*k, w*) values. (A-D) Expected density comparisons over fixed *k* or *w*. (E,F) Particular density comparisons over 1 million nucleotides from Chromosome X. (G,H) Expected density comparisons over *w* ∈ {*k, k* − 1}.

In the case *w* ∈ {*k, k* − 1} (Figures 2G,H), `GM` -`expected` generated minimizers that have expected density within 1% of the theoretical lower bound for forward schemes [12], which constitute a wide class of local schemes including all minimizers.

### 4.3. Density comparisons for popular HTS methods

To emphasize the relevance and potential impact of `GreedyMini`, we calculated the expected density factors of minimizers generated by `GM` -`expected` over (*k, w*) combinations used by popular HTS methods [23, 24, 25, 26, 27, 28, 29, 30, 31, 32]. We compared the results of `GreedyMini` to those of double decycling, miniception, open-closed syncmer, and random minimizer schemes.

For the combinations with *w > k* in four out of five cases, `GreedyMini` achieved the best result (Figure 3A, Supplementary Table S1). It loses to double decycling and open-closed syncmer schemes only for the pair (19, 30) used by GraphAligner. Among four considered combinations with *k > w*, `GreedyMini` shows the best result for three and second best (after open-closed syncmer) for the remaining one (Supplementary Table S1).

**Fig. 3.**
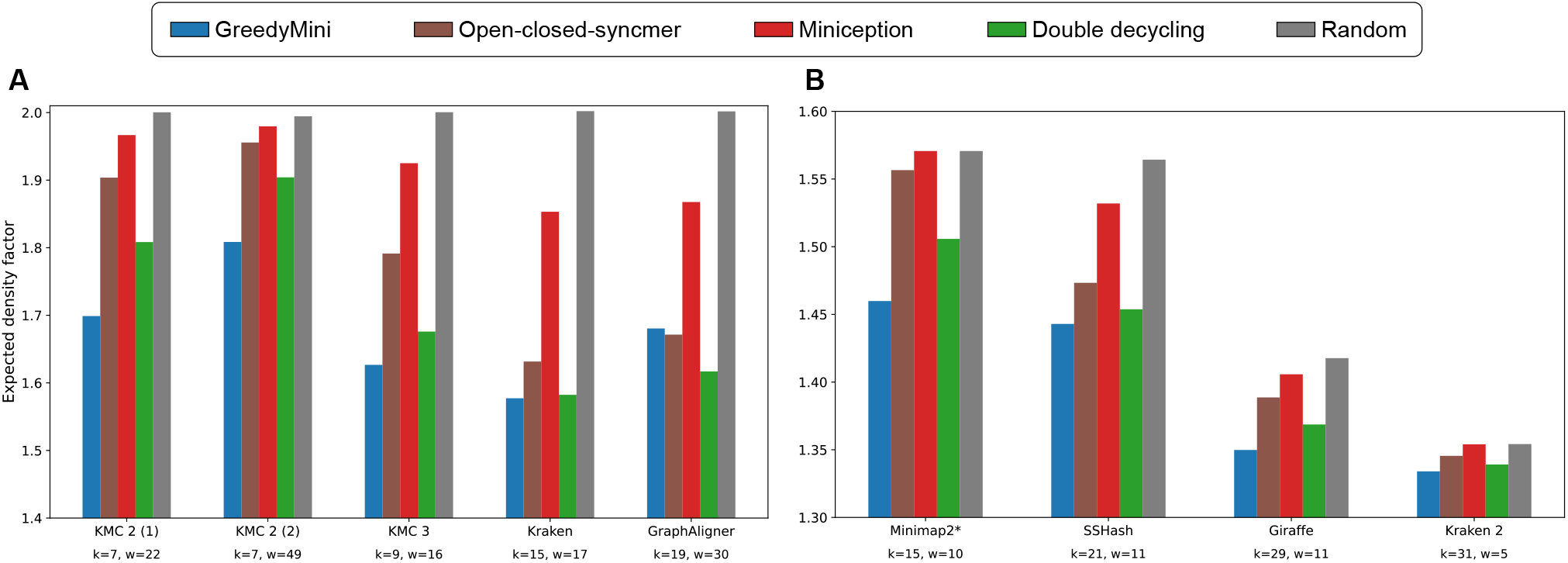
Density comparison over popular HTS methods. (A) Comparisons with other minimizers for *k < w*. (B) Comparison with other minimizers when plugged into a mod-minimizer scheme for *k > w*. *These (*k, w*) values also apply to Minimap [28] and MetaProb 2 [30]. We calculated the densities over a 10^7^-long random DNA sequences.

However, among all known forward schemes, the best results in the case *k > w* are achieved by the mod-minimizer scheme [15]. This scheme is not a minimizer, but uses a minimizer (for some *k*^*′*^ *< k*) as an intermediate step. Thus, we plugged the minimizers obtained by `GreedyMini` and all benchmarked methods into the mod-minimizer scheme, using the parameter *r* = 4 as in the original setting [20]. We measured the density of obtained schemes over a 10^7^-long random DNA sequence. In all four cases, `GreedyMini` showed the best result (Figure 3B, Supplementary Table S2).

### 4.4. *k*-mer rank-retrieval times

We next evaluated the sampling time for the DNA minimizers produced by `GreedyMini`. Sampling a string *S* consists of two tasks: retrieving the ranks of all *k*-mers and computing the minimum rank in a sliding window. The second task is independent of the minimizer used, so we focused on the rank-retrieval time.

We ran a code that assigns ranks to *k*-mers on a random DNA sequence of 3 · 10^9^ nucleotides, and calculated the average runtime over 10 runs for robust estimation. To simulate an order generated by `GM` -`expected` or `GM` -`particular`, we stored a binary order in memory and extended it by `Extend` -*σ*. For benchmarking, we used two ranking algorithms that do not require storing an order in memory: a trivial XOR-based hash function, which is used by some minimizers to create a pseudo-random order, and std::hash, which is the default hash implementation in C^++^. We ran our code on an AMD Ryzen™ 7 7800×3D (L1/L2/L3 cache size: 512KB/8MB/96MB) alongside 32GB of DDR5 RAM.

For values for which `GM` -`expected` generates DNA minimizers in a reasonable runtime (i.e., *k* ≤ 15), it is also very fast to retrieve *k*-mer ranks using the resulting minimizers (Figure 4). The rank-retrieval times for `GreedyMini` match those for the XOR-based hash function (less than 4 seconds over 3 billion nucleotides including the generation of the random input string). For 17 ≤ *k* ≤ 22, there was a small increase in time, which still remains well below the results of std::hash. Only for *k >* 22, the rank-retrieval time steeply increases, presumably switching to storing (some of) the order in the slowest cache or even in the main memory. Since any order generated by `GreedyMini` for *k >* 22 is likely to use a trivial *k*-extension from some smaller *k*^*′*^, then only the order of *k*^*′*^-mers needs to be stored. Thus, on the tested hardware the sampling time of the minimizers generated by `GreedyMini` is guaranteed to be competitive.

**Fig. 4.**
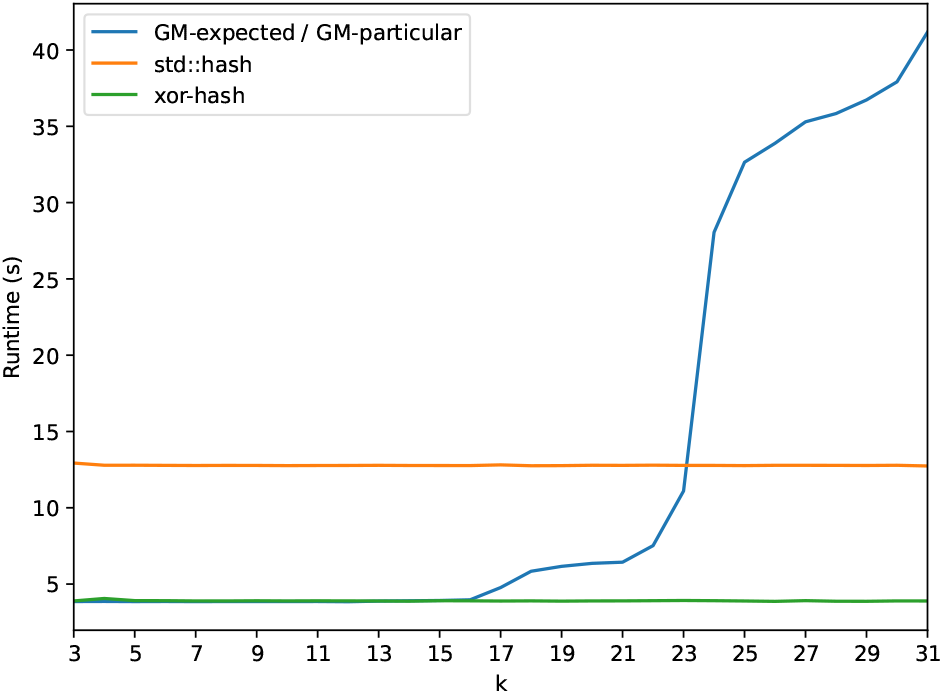
Runtime to assign ranks to *k*-mers over a 3 · 10^9^-long random DNA sequence, averaged over 10 runs.

### 4.5. Effect of individual steps on density results

We next measured and analyzed the impact of different steps in `GreedyMini` on the obtained densities. We tested the effect of transforming binary minimizers to DNA minimizers by `Extend` -*σ* and the effect of the local improvements by `Swap`.

The difference in expected density between the binary minimizers generated by stages Generate and Improve of `GM` -`expected` and the DNA minimizers obtained with `Extend` -*σ* in stage Extend was less than 1% for all minimizers generated over 5 ≤ *k* ≤ 15 and 3 ≤ *w* ≤ 15 (Supplementary Table S3); moreover, for *k* ≥ 9 this difference was well below 0.1% in all cases. Thus, we conclude that `Extend` -*σ* produces DNA minimizers with densities very similar to those of the original binary minimizers.

The contribution of `Swap` was minor for most (*k, w*) combinations we tested (with the orders generated by `GreedyE`). For most pairs (*k, w*) with 3 ≤ *w* ≤ 15, *k* ≤ 6, there was no reduction in expected density, possibly due to a global or local optimum. For 3 ≤ *w* ≤ 15 and 7 ≤ *k* ≤ 15, the average reduction in expected density was 0.26% ± 0.15% with the reduction 0.67% for the pair (13, 10) being the best improvement (Supplementary Table S4). In contrast, if the pair (*k, w*) is out of the range of `GreedyE`, the use of `Swap` in the `GM` -`improve` pipeline resulted in substantial improvements. For example, for the pair (12, 30) `Swap` has reduced the expected density by 4.35%.

Overall, we conclude that the principal role in constructing low-density minimizers is played by `GreedyE` /`GreedyP`, with mostly marginal effect of other steps on density.

### 4.6. Towards optimal minimizers

As already mentioned in Section 4.2, the densities of minimizers generated by `GreedyMini` pipelines are close to the lower bound on the density of forward schemes [12]. Moreover, the *binary* minimizers by `GreedyMini` pipelines in the process hit this lower bound for the following (*k, w*) pairs: (4, 3), (7, 3), (10, 3), (5, 4), and (6, 5). Apart from proving that, for some (*k, w*) combinations, minimizers are as good as general forward schemes, this result supports the conjecture that, within the range where `GreedyMini` is feasible to run, it generates minimizers with the density close to the lower bound. Based on this conjecture, we present in Figure 5 the landscape of density factors of our best binary minimizers as an approximation of such a landscape of minimum densities. The main features of this landscape are flat “highlands” (*w* ≥ *k* − 1), gradually ascending as *w* grows over *k*, bumpy “lowlands” (*w* ≤ *k* − 3), and a steep step between them (*k* − 3 ≤ *w* ≤ *k* − 1).

**Fig. 5.**
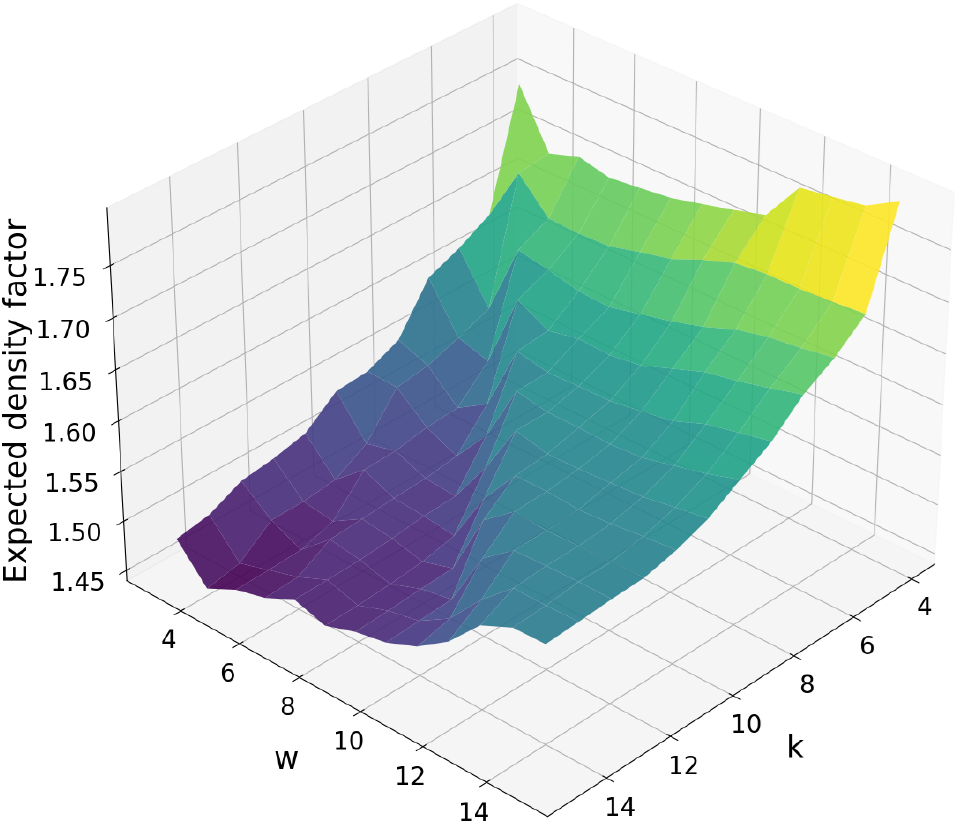
Density factors of the best binary minimizers generated by GreedyMini pipelines.

## 5. Discussion

In this paper, we tackled the problem of generating a minimizer that achieves low expected or particular density. We presented `GreedyMini`, a toolkit of novel methods to generate low-density minimizers, to improve a given minimizer, to extend to larger alphabet, *k*, and *w*, and to measure the density of a given minimizer. With these methods, we devised pipelines to generate low-density DNA minimizers for given (*k, w*) combinations, with the key role played by two variants of a greedy algorithm (`GreedyE` and `GreedyP` for expected and particular density, respectively).

We evaluated the performance of the minimizers generated by `GreedyMini` pipelines by calculating the achieved expected and particular densities, and conclude that `GreedyMini` pipelines generate minimizers that achieve the lowest density compared to all existing minimizer methods over a wide range of (*k, w*) combinations. This range includes the parameters used by popular HTS tools, such as Minimap2 [29], Kraken 2 [24], KMC 3 [26], and several others. Moreover, we observed that the achieved densities are very close to the theoretical lower bound in that range, and even hit the lower bound for forward schemes for the binary alphabet and several (*k, w*) pairs.

The main limitation of `GreedyMini` is the exponential dependence on *w* + *k* of the time needed to build a minimizer by `GreedyE`. Thanks to our novel minimizer-to-minimizer transformation to larger alphabets (`Extend` -*σ*), this dependence is only *O*(2^2*k*^ +*w*2^*w*+*k*^) per run of `GreedyE`. This is an acceptable construction time for a wide range of practical values of *k* and *w*. This range is further extended to larger *k* values due to `Extend` -*k* and to larger *w* values by `Extend` -*w* and by `GM` -`improve`. The use of `Extend` -*σ* reduces the exponential dependence of the space complexity in *k* to just *O*(2^*k*^), which is a feasible limitation. For *k* = 20, the lookup table occupies 4·2^20^ bytes (= 4MB), which surely fits in the cache memory, where an access operation takes *O*(1) time under many reasonable models. Indeed, we showed that the *k*-mer rank-retrieval time for such a lookup table is comparable to that of the computationally trivial XOR hash function. For larger *k*, one can generate a minimizer for *k* = 20 and use `Extend` -*k* multiple times, keeping the size of the lookup table within the same 4MB.

Our study raises several open questions. First, for which (*σ, k, w*) combinations minimizers are optimal among all forward schemes? Second, what is the complexity of the minimum-density problem, and can we generate minimum-density minimizers efficiently? Third, can we prove any guarantees for `GreedyE`, e.g. whether greedy is *always* better than random? Fourth, is there an algorithm that returns the rank of a *k*-mer in the order generated by `GM` -`expected` or `GM` -`particular` without storing the order explicitly?

There are several directions for future work. First, we plan to plug our new minimizers in schemes, such as strobemers [16], to improve the density and study the impact on other metrics, such as conservation. Second, we plan to extend `GreedyMini` to complete any partially constructed order to a UHS order, such as an order defined on a double-decycling set. Third, we plan to incorporate our new minimizers in existing HTS algorithms and data structures and study the impact on their performance in terms of runtime and memory usage.

## 6. Code availability

The toolkit and its source code are publicly available at: https://github.com/OrensteinLab/GreedyMini.

## 8. Author contributions statement

S.G., M.K, and A.S. performed the theoretical analysis. I.T. implemented and benchmarked the toolkit. S.G., A.S., and Y.O. led the project. All authors wrote the manuscript.

## 9. Acknowledgments

Shay Golan is supported by Israel Science Foundation grant no. 810/21. Ido Tziony acknowledges the cloud-computing credit support of the Israel Data Science and AI Initiative. Arseny Shur is supported by the ERC grant MPM no. 683064 under the EU’s Horizon 2020 Research and Innovation Programme and by the State of Israel through the Center for Absorption in Science of the Ministry of Aliyah and Immigration.

## Supplementary Notes

### S1. Preliminaries

#### S1.1. Strings as integers

In our algorithms, we view *k*-mers and explicitly stored windows as **integers written in** *σ***-ary** (padded with leading zeroes to the required length). As a result, many string operations, like extracting a *k*-mer in a given position, become unit-cost integer operations. In Algorithms S7 and S8, the integer operations are made explicit; the shift operation *υ* ≫_*σ*_ *n* deletes the last *n* digits (characters) of *υ*.

#### S1.2. Tries and reversed tries

Given a set *U* of strings, its *trie* (or *prefix tree*) *T*_*U*_ is defined as follows. The set of nodes is the set of all prefixes of the strings in *U*; the set of edges is the set of all ordered pairs (*z, za*) of nodes, where *a* ∈ Σ. Similarly, the *reversed trie* 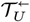 has all *suffixes* of the strings in *U* as nodes and all ordered pairs of nodes of the form (*z, az*) as edges.

We use tries and reversed tries of sets of (*k* + *w*)-windows, computing node characteristics during a depth-first search (DFS) without explicitly storing the whole tree. We assume that each node *z* of a trie (respectively, reversed trie) with |*z*| ≥ *k* is labeled by its *k*-suffix (respectively, *k*-prefix).

#### S1.3. Relation between proofs of theorems

The algorithms proving Theorems 1, 2, and 5 share common ideas. We present these ideas from the simplest to the hardest case, so we first prove Theorem 5, then Theorem 2, and finally Theorem 1. After that, we prove Theorems 3 and 4, which also share common ideas in the proofs.

### S2. Proof of Theorem 5

To calculate the density *d*_(*ρ,w*)_, we count the (*w*+*k*)-windows charged by *ρ* and divide the result by *σ*^*w*+*k*^. We focus on counting charged windows, doing a separate count for windows that are *prefix-charged* (the minimal *k*-mer is a prefix) and *suffix-charged* (the minimal *k*-mer is a unique suffix). Two different approaches, described below, result in the algorithms `DenDFS` and `DenDP`, reaching the complexity stated in Theorem 5 (a) and (b), respectively.

#### S2.1. *Algorithm* DenDFS

Let *U* be the set of all windows with some *k*-prefix *u*. The trie *T*_*U*_ of *U* consists of the path from the root to *u* and a complete *σ*-ary tree of depth *w*, rooted at *u*. In order to count prefix-charged windows with the prefix *u*, we run a recursive DFS on this complete subtree by calling `DFSp` (*u, u*, 0, `rank` _*ρ*_(*u*)) (Algorithm S1).

With each node *z* we associate the minimum rank *r*_*z*_ of a *k*-mer occurring in *z*, i.e., *r*_*z*_ = min{`rank` _*ρ*_(*u*^*′*^) | *u*^*′*^ occurs in *z*}. If *z* is internal, it recursively calls all its children; the minimum rank for a child *za* is computed as min{*r*_*z*_, `rank` _*ρ*_(*z*_*a*_)}, where *z*_*a*_ is the *k*-suffix of *za*. After that, *z* returns the number of prefix-charged windows in its subtree, summing up the values returned by its children. If *z* is a leaf, i.e., a window in *U*, it returns 1 if it is prefix-charged (which means exactly *r*_*z*_ = `rank` _*ρ*_(*u*)) and 0 otherwise.

##### Algorithm S1: function DFSp for recursive count of prefix-charged (*w*+*k*)-windows

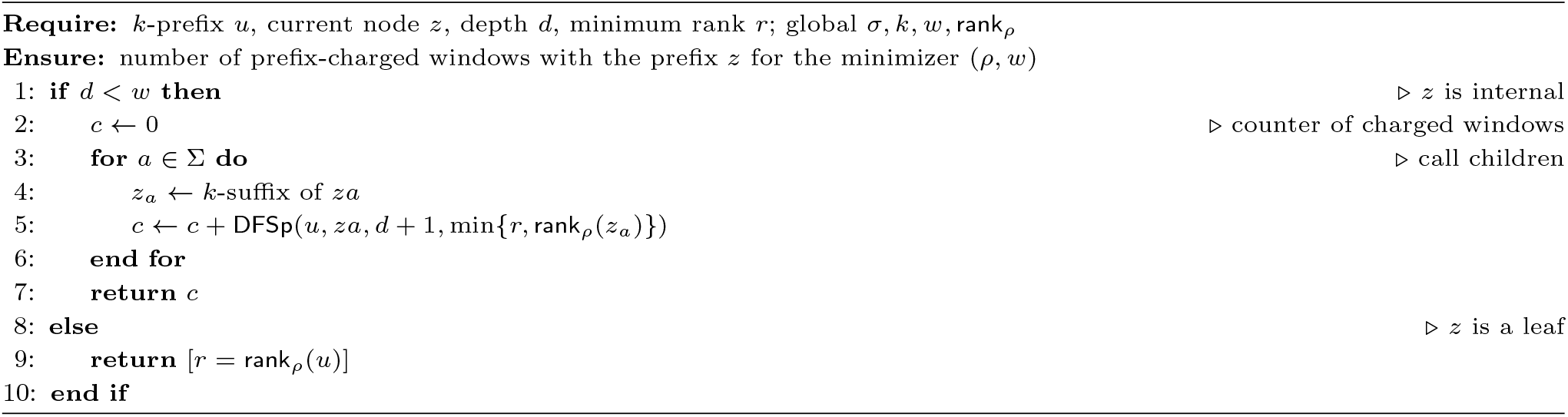

The call `DFSp` (*u, u*, 0, `rank` _*ρ*_(*u*)) spends *O*(*σ*) time per internal node of 𝒯_*U*_ and *O*(1) time per leaf, which is *O*(*σ*^*w*^) in total. It stores *O*(1) numbers per active recursive call, which means *O*(*w*) space in total.

We count suffix-charged windows with the suffix *u* in a symmetric way, using a recursive function `DFSs` on the reversed trie 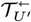, where *U*^*′*^ consists of all (*w* + *k*)-windows with the suffix *u*. The difference with `DFSp` is that a leaf *z* returns 1 if *z* contains, apart from the suffix *u*, no *k*-mers of rank smaller than or **equal to** `rank` _*ρ*_(*u*). To ensure correct processing, we set the value returned by a leaf *z* to [*r*_*z*_ = `rank` _*ρ*_(*u*) + 1] and start the search by calling `DFSs` (*u, u*, 0, `rank` _*ρ*_(*u*) + 1). This call obviously satisfies the complexity bounds computed above for `DFSp`.

Algorithm `DenDFS` calls `DFSp` (*u, u*, 0, `rank` _*ρ*_(*u*)) and `DFSs` (*u, u*, 0, `rank` _*ρ*_(*u*) + 1) for each ranked *k*-mer *u* and sums up the results to get the number of windows charged by *ρ*. The algorithm spends *O*(|*H*|*σ*^*w*^) time and *O*(*w*) space, as required by statement (a).

#### S2.2. Algorithm DenDP

Recall that *order-k deBrujin graph over* Σ is a directed *σ*-regular graph, having all *σ*-ary *k*-mers as nodes and all pairs (*au, ub*), where *u* ∈ Σ^*k*−1^, *a, b* ∈ Σ, as edges. If an edge (*au, ub*) is labeled by *b*, the graph becomes a deterministic finite automaton *B*. We view ℬ as a *transition table* with rows indexed by *k*-mers and columns indexed by letters; the entry ℬ [*u, a*] contains the successor of *u* by the letter *a*.

We complete the UHS order *ρ* to a linear order 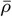, assigning the remaining ranks lexicographically. Then we replace all elements and all row indices in *B* with their 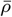-ranks, and sort the rows. The resulting table is referred to as ℬ_*ρ*_.

To count (*w* + *k*)-windows charged due to some *k*-mer *u*, we proceed by dynamic programming. Let `pref` _*r*_ be a two-dimensional table such that `pref` _*r*_ [*i, j*] is the number of strings of length *k* + *j* having the *k*-prefix of rank *r*, the *k*-suffix of rank *i*, and all other *k*-mers of rank at least *r*. Then the number of (*w* + *k*)-windows prefix-charged due to rank-*r k*-mer is Σ*i* ≥*r* `pref` _*r*_ [*i, w*]. Similarly, let `suff` _*r*_ be a two-dimensional table such that `suff` _*r*_[*i, j*] is the number of strings of length *k* + *j* having the *k*-suffix of rank *i* and all other *k*-mers of rank greater than *r*. Then the number of (*w* + *k*)-windows suffix-charged due to rank-*r k*-mer is suff_*r*_[*r, w*].

Note that `pref` _*r*_ [*i*, 0] = [*i* = *r*], `suff` _*r*_[*i*, 0] = 1, and the DP rules for `pref` _*r*_ and `suff` _*r*_ are almost the same. Namely, `pref` _*r*_ [*i, j* + 1] =Σ_𝓁_ `pref` _*r*_ [𝓁, *j*], where the summation is over all ranks 𝓁 ≥ *r* such that the *k*-mer of rank *i* is a successor of the *k*-mer of rank 𝓁. The rule is computed by the function `DPp` (Algorithm S2), which uses the transition table ℬ_*ρ*_ to propagate the counts from the *j*’th column to the (*j* + 1)th column along the edges of the deBrujin graph. The DP rule for suffixes looks the same: `suff` _*r*_[*i, j* + 1] =Σ_𝓁_ `suff` _*r*_[𝓁, *j*]; the only difference is that the inequality for 𝓁 is strict: 𝓁 *> r*. This rule is computed by the function `DPs` (Algorithm S3).

##### Algorithm S2: function DPp for count of prefix-charged (*w*+*k*)-windows with a prefix of a given rank

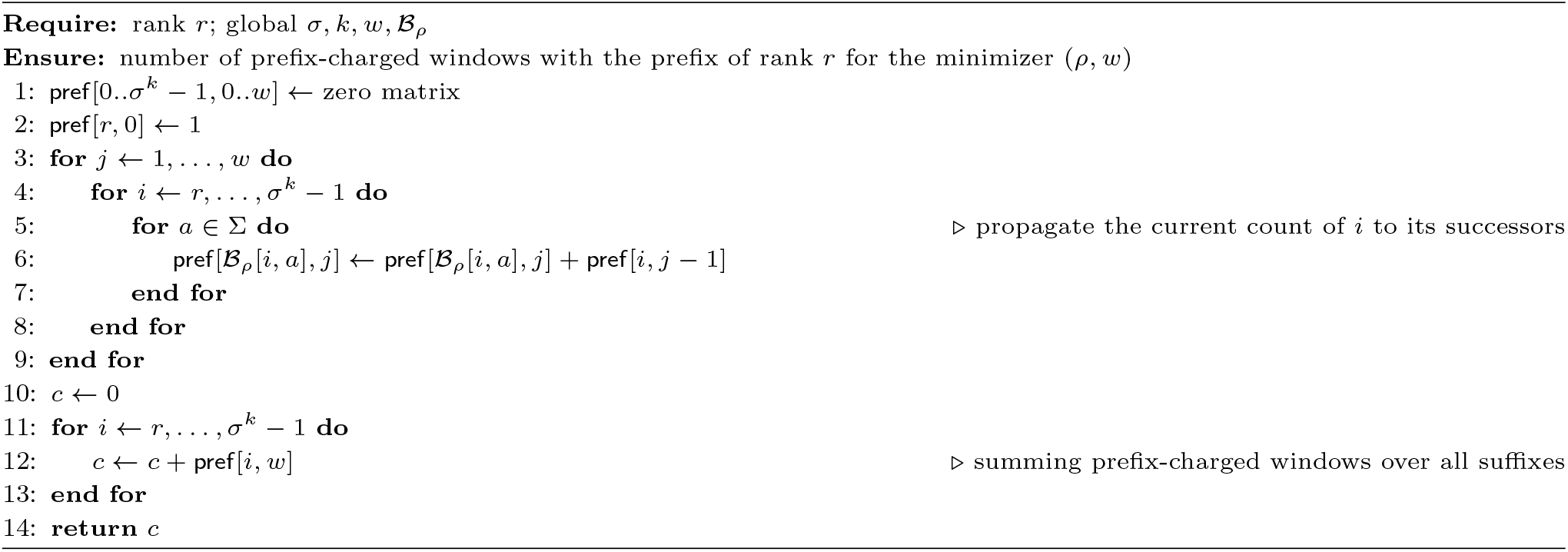

##### Algorithm S3: function DPs for count of suffix-charged (*w*+*k*)-windows with a suffix of a given rank

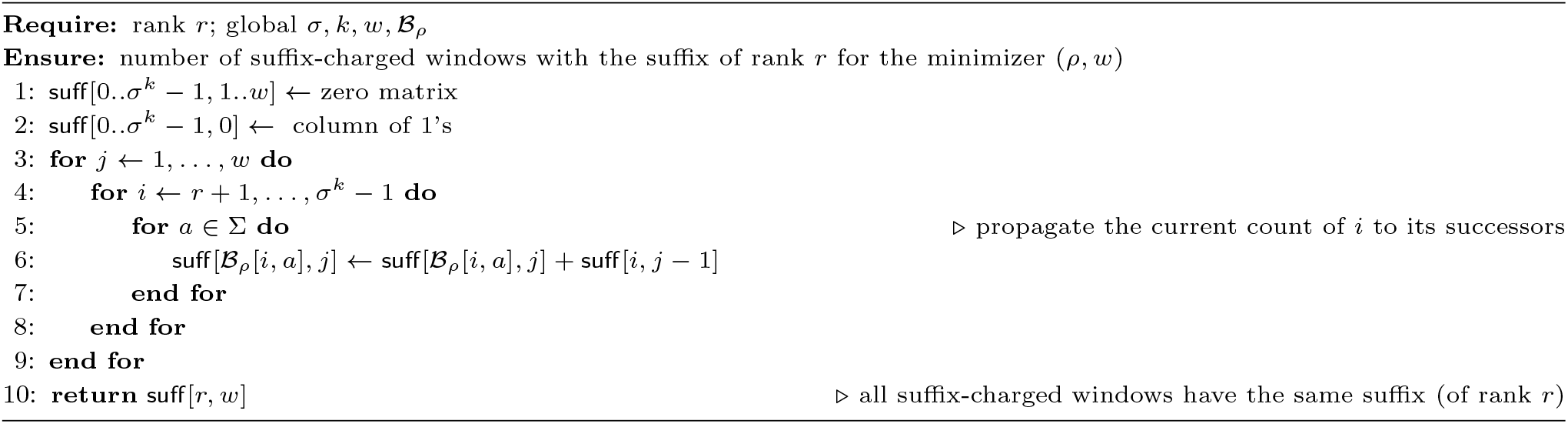

Algorithm S2 works in *O*(*wσ*^*k*^) time (recall that we assume *σ* to be a constant) and uses *O*(*σ*^*k*^) space, as just two columns of the DP table are stored at every moment. The same bounds apply for counting suffix-charged windows.

The algorithm `DenDP` creates the table ℬ_*ρ*_ and then calls `DPp` (*r*) and `DPs` (*r*) for each rank *r* assigned in *ρ*. As ℬ_*ρ*_ can be trivially computed from `rank` _*ρ*_ in *O*(*σ*^*k*^) time and space, `DenDP` works in *O*(|*H*|*wσ*^*k*^) time and *O*(*σ*^*k*^) space, as required in statement (b). Theorem 5 is proved.

##### Remark S1

*An additional feature of the dynamic programming algorithm* `DenDP` *is that while computing d*_(*ρ,w*)_ *it can, within the same time and space bounds, report the densities of* ***all*** *minimizers* (*ρ, w*^*′*^) *with w*^*′*^ ≤ *w*.

### S3. Proof of Theorem 2

Let *ρ* be a UHS order for *w, H* be its UHS, and *r* be a rank such that both *r* and *r*+1 are assigned in *ρ*. Let *ρ*^*′*^ be the order obtained from *ρ* by swapping *k*-mers *u* and *u*^*′*^ such that `rank` _*ρ*_[*u*] = *r*, `rank` _*ρ*_[*u*^*′*^] = *r* + 1. We define the *cost function* 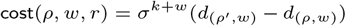, equal to the difference between the numbers of (*w* + *k*)-windows charged by *ρ*^*′*^ and by *ρ*. We focus on the efficient computation of `cost` (*ρ, w, r*), since all other operations in the function `Swap` are straightforward (Algorithm S4). Similar to Theorem 5, we present two algorithms computing the cost function: the DFS-based algorithm and the DP-based algorithm are used to prove the statements (a) and (b) of Theorem 2, respectively. Both algorithms analyse the difference between *ρ* and *ρ*^*′*^, avoiding direct computation of 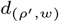.

#### Algorithm S4: function Swap for swapping two *k*-mers with consecutive ranks

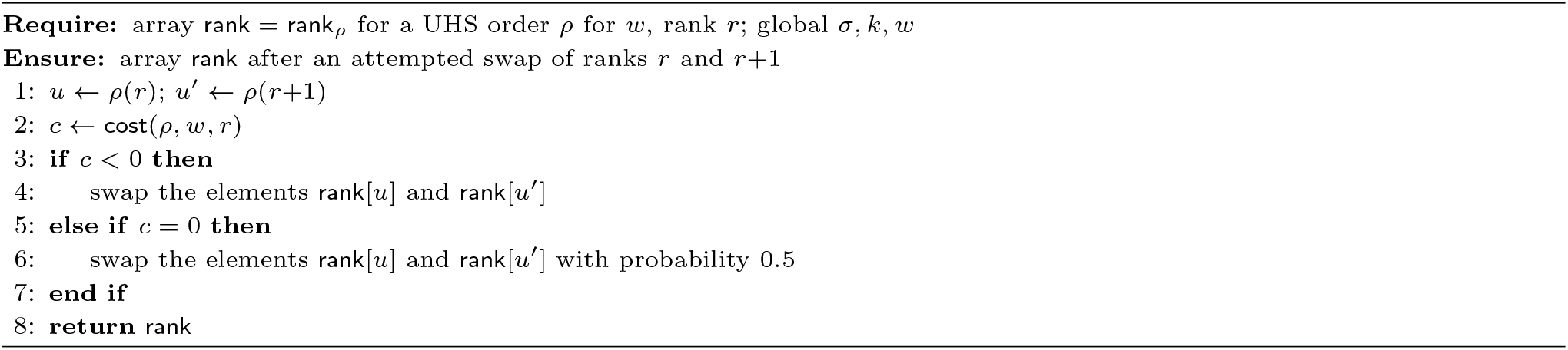

#### S3.1. *Algorithm* SwapDFS

In this section, compute `cost` (*ρ, w, r*) by considering only those (*w*+*k*)-windows that could have different status (free/charged) for *ρ* and *ρ*^*′*^. If a window *v* changed its status, then its minimum-rank *k*-mer in *ρ* is *u* and its minimum-rank *k*-mer in *ρ*^*′*^ is *u*^*′*^. Hence *v* contains both *u* and *u*^*′*^; moreover, either *u* or *u*^*′*^ is a prefix and/or a suffix of *v*. Therefore, it is sufficient to check four sets of windows: those with prefix *u*, prefix *u*^*′*^, suffix *u*, and suffix *u*^*′*^. The number of windows in these sets is *O*(*σ*^*w*^).

Let *U* be the set of all windows with the prefix *u* and let *v* ∈ *U*. If *v* contains a *k*-mer of rank *< r*, then it has the same status for both *ρ* and *ρ*^*′*^. Otherwise, it is charged by *ρ*; this charged window is free for *ρ*^*′*^ if it contains *u*^*′*^, but not as a unique suffix. Respectively, it suffices to count the windows that are charged by *ρ* and free for *ρ*^*′*^, as the opposite change of status is impossible. Similar to the proof of Theorem 5, we make use of the trie 𝒯_*U*_ of *U*. It consists of a path from the root to *u*, followed by a complete *σ*-ary tree of depth *w*. We additionally assign to each internal node *z* a flag indicating whether *u*^*′*^ occurs in *z* and run a DFS on the complete subtree, calling the function `pr_cost` (`rank` _*ρ*_, *r, u, u*^*′*^, *u*, 0, `False`) (Algorithm S5).

During the DFS, each node returns the number of (*w* + *k*)-windows in its subtree that are charged by *ρ* and free for *ρ*^*′*^. For a node *z* with the label *z*^*′*^, if `rank` [*z*^*′*^] *< r*, we return 0. Otherwise, the processing depends on the type of the node. Let *z* be internal. If *z*^*′*^ = *u*^*′*^, we set the flag; otherwise, we copy its value from the parent of *z*. After that, we recursively call all children of *z* and return the sum of the values they returned. Now let *z* be a leaf. It returns 1 if the flag of its parent is set and 0 otherwise. Clearly, the root *u* of the subtree returns the required value. The processing time is *O*(*σ*^*w*^) and the space is *O*(*w*), as we store only *O*(1) numbers per each of *O*(*w*) active recursive calls. We then process the set *U*^*′*^ of windows with the prefix *u*^*′*^ in the same way as *U*, calling `pr_cost` (`rank` _*ρ*_, *r, u*^*′*^, *u, u*^*′*^, 0, `False`) to count the windows that are charged by *ρ*^*′*^ and free for *ρ*.

##### Algorithm S5: function pr_cost for recursive count of status-changing *k*-mers

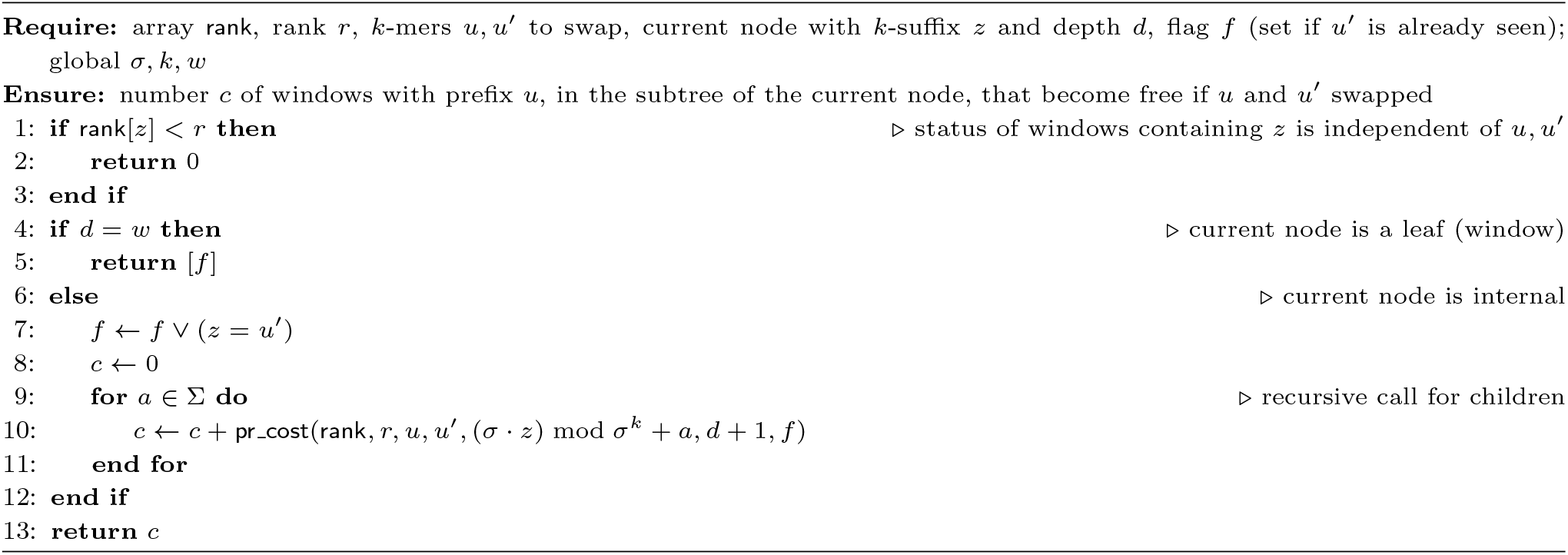

Consider the set *Ū* of the windows with the suffix *u*. The options for *v* ∈ *Ū* are the following. If *v* contains a *k*-mer of rank *< r* or it does not contain *u*^*′*^, then it has the same status in *ρ* and *ρ*^*′*^. If *v* has the prefix *u* or *u*^*′*^, we ignore it to avoid double count. Otherwise, *v* is free for *ρ*^*′*^ and charged by *ρ* if and only if it has a single occurrence of *u*. We process *Ū*similar to *U*, counting the windows that are charged by *ρ* and free for *ρ*^*′*^, but in the **reversed** trie 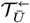. We use a DFS over the complete subtree of 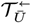 rooted at *u* and adjust the processing of a node according to the conditions listed above, getting the function `sf_cost` (Algorithm S6). It has the same space and time bounds as `pr_cost`, and the set *Ū*^*′*^ of the windows with the suffix *u*^*′*^ is processed in exactly the same way as *Ū*. Finally, `cost` (*ρ, w, r*) = `pr_cost` (`rank` _*ρ*_, *r, u*^*′*^, *u, u*^*′*^, 0, `False`) + `sf_cost` (`rank` _*ρ*_, *r, u*^*′*^, *u, u*^*′*^, 0, `False`) − `pr_cost` (`rank` _*ρ*_, *r, u, u*^*′*^, *u*, 0, `False`) − `sf_cost` (`rank` _*ρ*_, *r, u, u*^*′*^, *u*, 0, `False`) is computed in *O*(*σ*^*w*^) time and *O*(*w*) space. Adding *O*(*σ*^*k*^) time for finding the *k*-mers *u* and *u*^*′*^ (Algorithm S4, line 1), we get the complexity bounds of Theorem 2(a).

##### Algorithm S6: function sf_cost for recursive count of status-changing *k*-mers

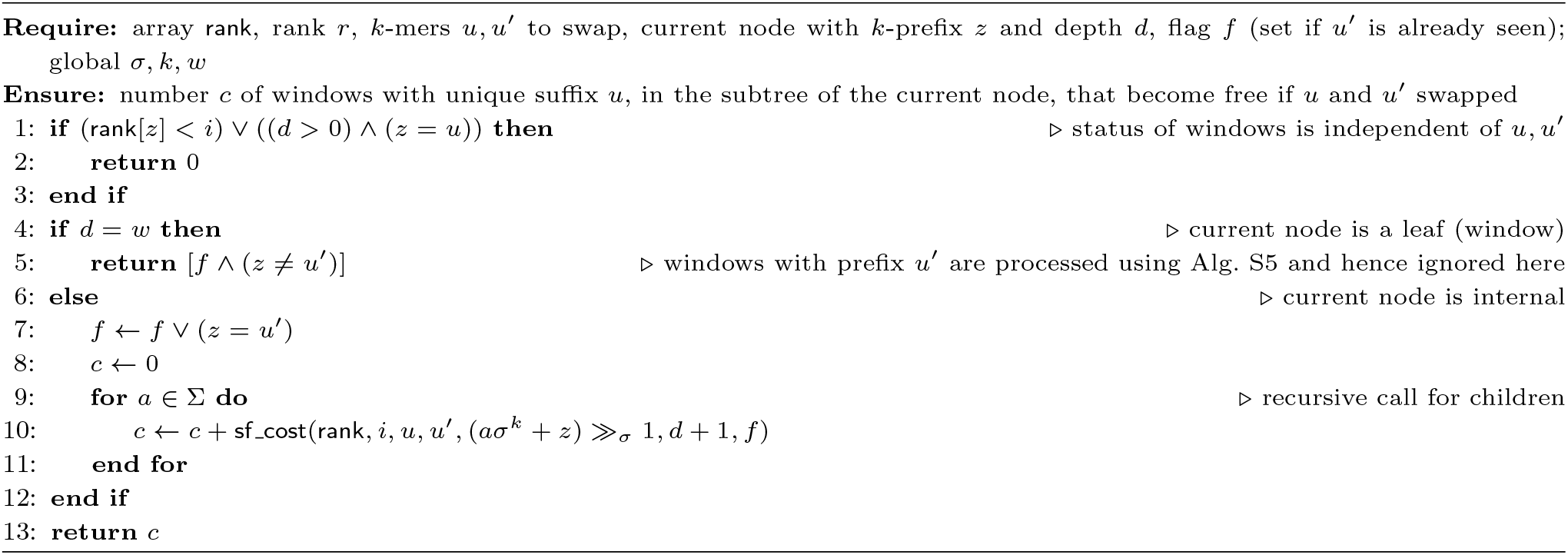

Algorithm `SwapDFS` runs multiple iterations of local search. Each iteration calls `Swap` (*ρ, w, r*) for the current UHS order *ρ* and a randomly chosen rank *r*; within this call, the cost function is computed by the depth-first search as described above.

#### S3.2. Algorithm SwapDP

From Section S2, we use the function `DPp` counting (*w* + *k*)-windows prefix-charged due to a *k*-mer of a given rank (Algorithm S2), and its counterpart `DPs` for suffix-charged windows (Algorithm S3). Let *c*(*ρ, w, i*) be the number of (*w* + *k*)-windows charged by *ρ* due to the rank-*i k*-mer. Then `cost` (*ρ, w, r*) = *c*(*ρ*^*′*^, *w, r*) + *c*(*ρ*^*′*^, *w, r* + 1) − *c*(*ρ, w, r*) − *c*(*ρ, w, r* + 1). Accordingly, we compute *c*(*ρ, w, r*) = `DPp` (*r*) + `DPs` (*r*) and *c*(*ρ, w, r* + 1) = `DPp` (*r* + 1) + `DPs` (*r* + 1) using the transition table ℬ_*ρ*_, then edit ℬ_*ρ*_ to get 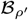, and compute *c*(*ρ*^*′*^, *w, r*), *c*(*ρ*^*′*^, *w, r* + 1) in the same way, but with the table 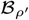. Note that given ℬ_*ρ*_, *ρ*, and `rank` _*ρ*_, the transition table 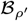 can be computed in constant time. Indeed, we take *u* = *ρ*(*r*), compute all its predecessors by *O*(1) arithmetic operations, extract their ranks from `rank` _*ρ*_ and replace *r* with *r* + 1 in the rows of ℬ_*ρ*_ corresponding to these ranks. Then we do the same with *u*^*′*^ = *ρ*(*r* + 1), and replace *r* + 1 with *r* in the obtained rows. Finally we swap the rows *r* and *r* + 1, getting 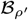.

In this way, `cost` (*ρ, w, r*) is computed within the complexity bounds of Algorithm S2, i.e., in *O*(*wσ*^*k*^) time and *O*(*σ*^*k*^) space. Then we immediately have statement (b) of Theorem 2.

Algorithm `SwapDP` runs multiple iterations of local search. Before the first iteration, it spends *O*(*σ*^*k*^) time to compute 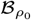 for the input UHS order *ρ*_0_. Each iteration calls `Swap` (*ρ, w, r*) for the current UHS order *ρ* and a randomly chosen rank *r*; within this call, the cost function is computed by the dynamic programming algorithms `DPp` and `DPs` as described above. The changes in the transition table made during this computation are retained if the swap was successful and reversed otherwise.

### S4. Proof of Theorem 1

As with Theorems 2 and 5, both DFS-based and DP-based solutions are possible. For an unranked *k*-mer *u*, counting `score` (*u*) = |*Y*_*u*_|*/*|*X*_*u*_| can be done in *O*(*w* ^*k*^) time by variants of the functions `DPp` (Algorithm S2) and `DPs` (Algorithm S3). However, this means *O*(*wσ*^2*k*^) time per iteration, which is rather inefficient. Because of this, we stick to the DFS solution.

We maintain arrays `X`, `Y`, and `rank`, each of size *σ*^*k*^. For every *k*-mer *u*, the counters `X` [*u*] and `Y` [*u*] store the current values of |*X*_*u*_| and |*Y*_*u*_|, respectively; `rank` is used to store the assigned ranks over all *k*-mers. To initialize `X` and `Y` it suffices to loop through all windows, incrementing the counters of all *k*-mers occurring in the current window. This can be done in *O*(*wσ*^*w*+*k*^) time in a trivial way. Then at most *σ*^*k*^ iterations follow, each consisting of two steps: choosing the *k*-mer to assign rank to and updating the arrays `X`, `Y` (for convenience, the rank is assigned *after* the updates). Computing the pool of low-scored *k*-mers and choosing a random *k*-mer from it can be done in time proportional to the size of `X` and `Y`, i.e., *O*(*σ*^*k*^).

Assume that the *k*-mer *u* is chosen and consider the update step (Algorithm S7). For every *i* ∈ [0, *w*+1), exactly *σ*^*w*^ windows contain *u* at position *i*. Hence, the number of windows containing *u* is *O*(*wσ*^*w*^), and to fit into the time bound we need to spend amortized *O*(1) time per such window while skipping all windows that do not contain *u*. We group windows containing *u* in *u-blocks*. A *u*-block is a set of all windows *v* sharing the same prefix *pu*, where *p* is followed by the leftmost *u* in *v*. The number of *u*-blocks with the leftmost *u* starting at position *i* equals the number of options for *p*, which is at most *σ*^*i*^. Hence, the total number of *u*-blocks is at most 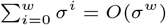.

#### Algorithm S7: function update_counts

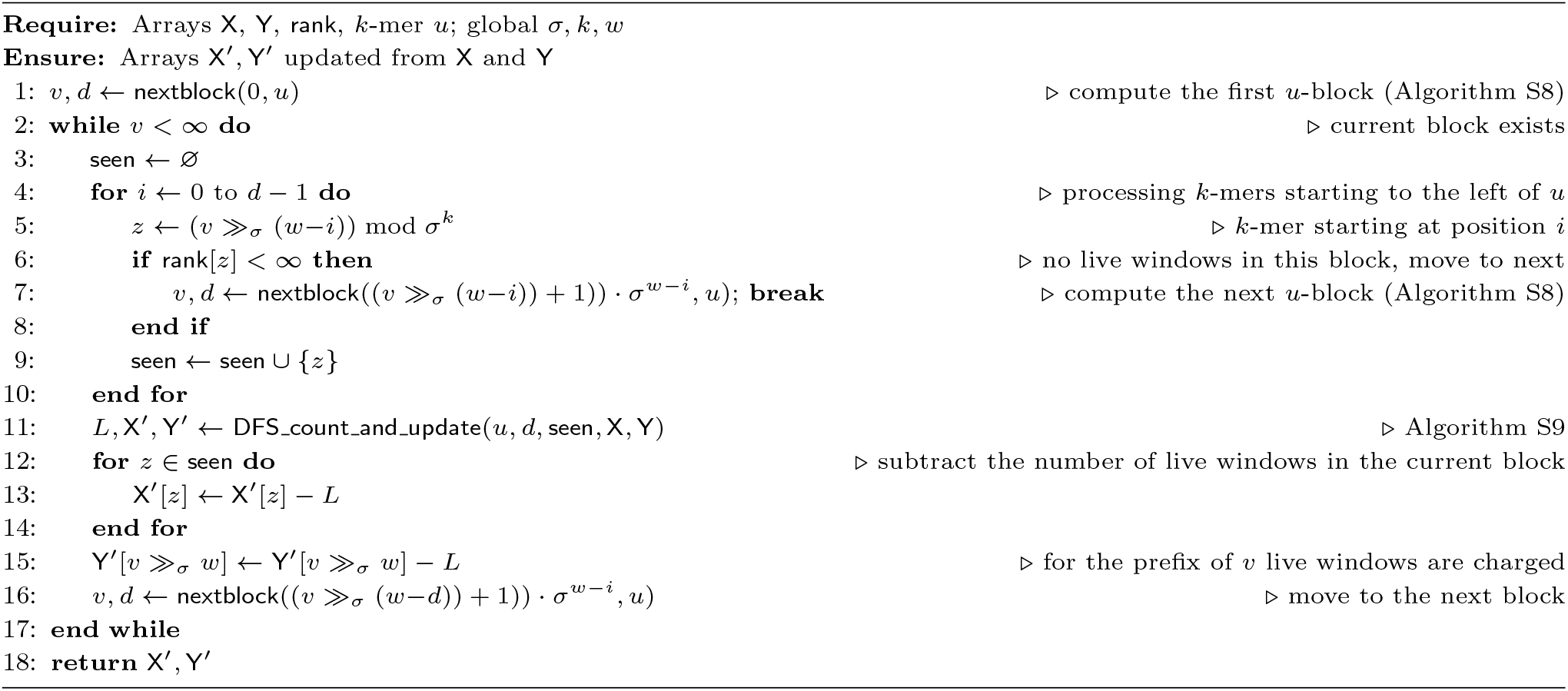

We process one block at a time, navigating between blocks with the help of an auxiliary function `nextblock` (*υ, u*) (Algorithm S8): it returns the smallest window *υ*^*′*^ that is greater than the window *υ* and contains *u*, or `null` if no such window exists. In other words, `nextblock` (*υ, u*) returns the first window of the next *u*-block. To implement `nextblock` (*υ, u*), we compute, for every *i* ∈ [0, *w*+1), the smallest window *υ*_*i*_ (if any) that is greater than *υ* and contains *u* at position *i*. Computing *υ*_*i*_ from *υ, u, i* requires a constant number of arithmetic operations. Hence, `nextblock` (*υ, u*) can be computed in *O*(*w*) time.

#### Algorithm S8: function nextblock

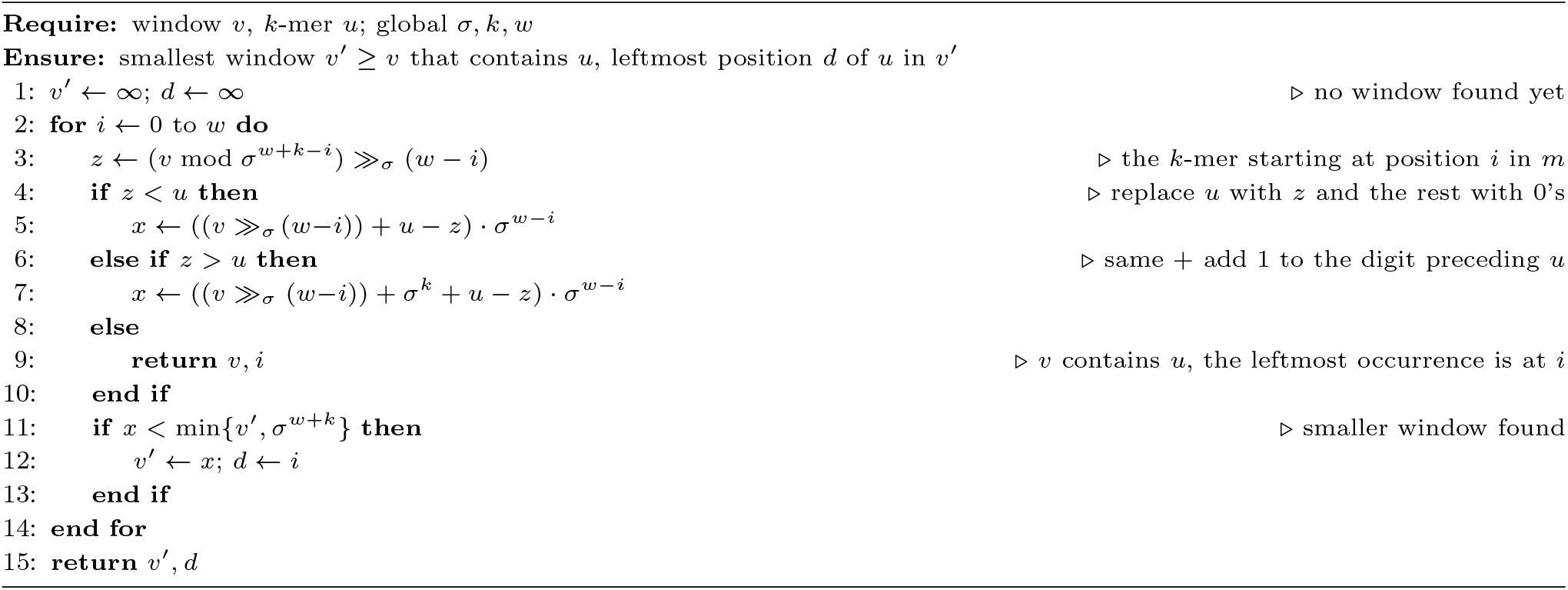

Consider processing a *u*-block *U* (one iteration of the **while** cycle in Algorithm S7). The goal is to count, for each unranked *k*-mer *u*^*′*^ occurring in at least one window of *U*, (i) the number of live windows from *U* containing *u*^*′*^ and (ii) the number of live windows from *U* containing *u*^*′*^ as a prefix or as a unique suffix. These numbers are then subtracted from `X` [*u*^*′*^] and `Y` [*u*^*′*^] respectively; in particular, `X` [*u*] and `Y` [*u*] become zeroes. Let *pu* be the common prefix of all windows in *U*, *i* = |*p*|, and let 𝒯_*U*_ be the trie of *U*. Each node of 𝒯_*U*_ is identified with a prefix of some windows from *U*, and leaves are identified with the windows themselves. The trie consists of a single path from the root to the node *pu* followed by a complete *σ*-ary tree of depth *w* − *i* below *pu*. Recall that each node of this complete tree is labeled with its *k*-suffix.

We first check all *k*-mers in *pu* except for *u*; if any of them has an assigned rank, then *U* contains no live windows, so the processing is finished (since all numbers to be subtracted from the cells of `X` and `Y` are zeroes). Otherwise, we put all of them in a set `seen` of occurred *k*-mers, associate `seen` with the node *pu* of 𝒯_*U*_, and run a recursive DFS (Algorithm S9) on the complete subtree of 𝒯_*U*_.

#### Algorithm S9: function DFS count and update

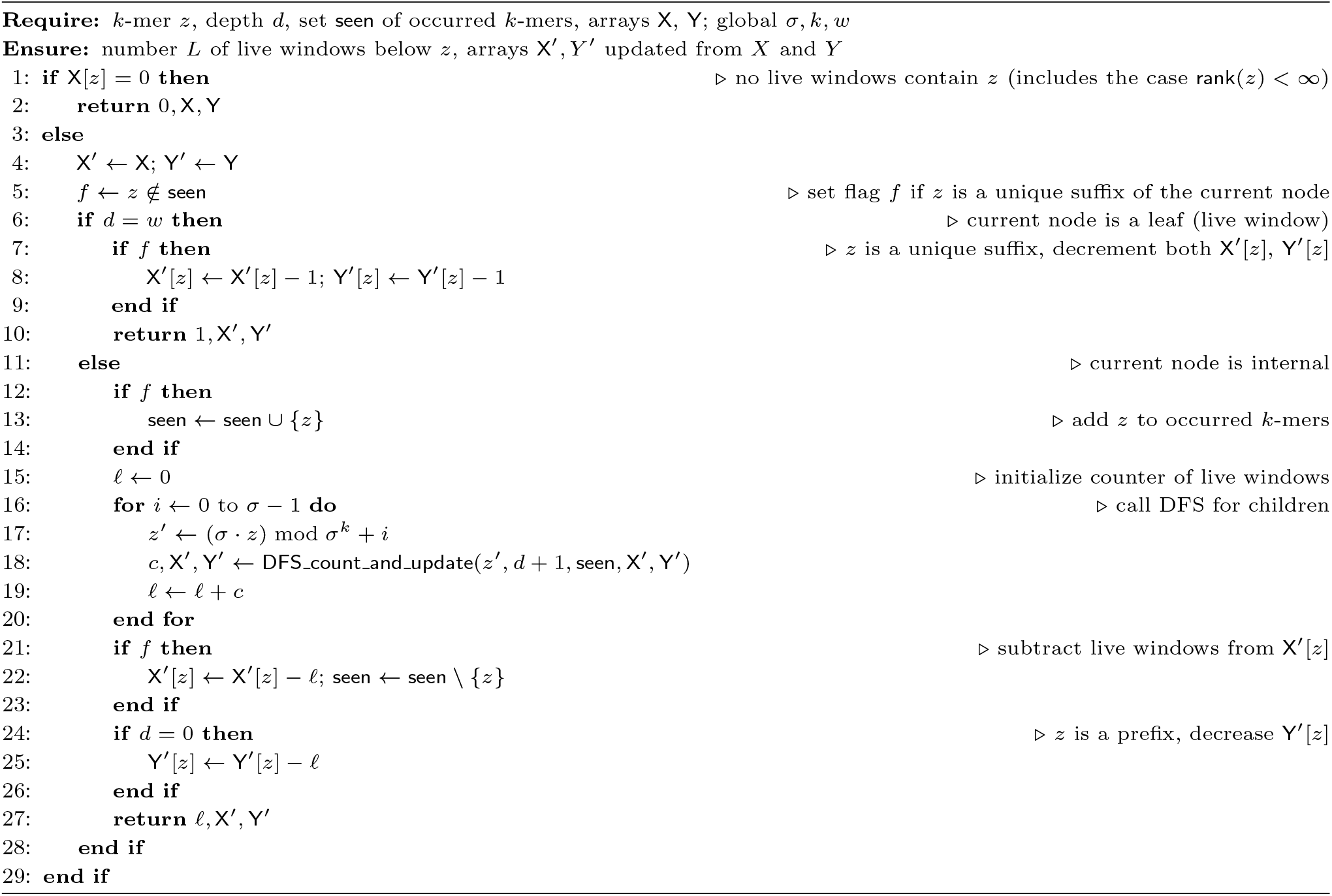

When the DFS reaches a node *z* with the label *z*^*′*^, we first check if *z*^*′*^ has an assigned rank. If yes, there are no live windows in the subtree of *z*, so this subtree is skipped and *z* returns 0. Otherwise, we set a boolean flag if *z*^*′*^ ∈ `seen`. After that, if *z* is an internal node, we insert *z*^*′*^ into `seen` if it was not there, and recursively call the children of *z*. When all children return values, we compute and return their sum *s*. If the boolean flag was not set, then before returning we delete *z*^*′*^ from `seen` and subtract *s* from `X` [*z*^*′*^] (if the flag *was set*, the subtraction will be done when processing the topmost occurrence of *z*^*′*^ among ancestors of *z*). Finally, if *z* is a leaf, we just return 1; if the flag was not set, before returning we subtract 1 from both `X` [*z*^*′*^] and `Y` [*z*^*′*^], since *z*^*′*^ is a unique suffix of a live window *z*.

When the DFS is finished, the root *pu* of the complete subtree returns the number ŝ of live windows in *U*. Now it remains to subtract ŝ from `X` [*u*^*′*^] for each *k*-mer *u*^*′*^ in *pu* (excluding *u*, which is already processed) and also from `Y` [*u*^*′′*^] for the *k*-prefix *u*^*′′*^ of *pu*. Thus, we reached the claimed goal.

When all *u*-blocks are processed, each value `X` [*u*^*′*^] (respectively, `Y` [*u*^*′*^]) was decreased by the number of live windows that contain *u* and also *u*^*′*^ (respectively, also *u*^*′*^ as a prefix or as a unique suffix). Since the live windows containing *u* are exactly those losing the state of “live” after the current iteration, the arrays `X` and `Y` are updated correctly.

It remains to check the time and space complexity. Processing a *u*-block *U* requires *O*(*w*) time for the *k*-mers in the common prefix *pu* and *O*(1) time per node of the complete subtree of the prefix tree 𝒯_*U*_. As there are *O*(*σ*^*w*^) *u*-blocks and the size of a complete subtree is proportional to the number of leaves in it, each of two processing times sums up to *O*(*wσ*^*w*^) over all *u*-blocks.

We also make *O*(*σ*^*w*^) calls to the `nextblock` function to move to the next *u*-block after finishing with the current one; as one call costs *O*(*w*) time, we arrive at the same bound. Adding *O*(*σ*^*k*^) time to select the *k*-mer *u* and multiplying the result by the number of iterations, we get the claimed time bound.

As for the space complexity, we do not store the subtree in which we run the DFS: the navigation is by arithmetic operations, so at any moment we store the set `seen` of *O*(*w*) size plus *O*(1) numbers per each of *O*(*w*) active calls. Adding the size of the arrays `X`, `Y`, and `rank`, we get the desired space bound. Theorem 1 is proved.

### S5. Discussion on *σ*- and *k*-extensions

In this section we analyze whether the straightforward density-preserving *σ*- and *k*-extensions of a minimizer are minimizers.

#### S5.1. σ-extension

Let *f* = (*ρ, w*) be a minimizer with the parameters (*σ, k, w*) and *γ >* 1 be an integer. Let 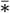 be any *projection* function from the alphabet {0,…, *σγ* − 1} onto {0,…, *σ* − 1}. The local scheme *f*_*γ*_ with the parameters (*σγ, k, w*) is defined as follows: in every window *υ, f*_*γ*_ picks the position chosen by *f* in the *projection* 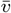 of *υ*, obtained from *v* by applying 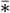 to each letter of *υ*.

##### Lemma 2

*f*_*γ*_ is not a minimizer.

*Proof* Suppose that *f*_*γ*_ is a minimizer (*π, w*). By pigeonhole principle there must be a symbol *s* ∈ {0,…, *σ* − 1} such that for two distinct *i, j* ∈ {0,…, *σγ* − 1} we have 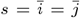. Consider the *k*-mer *u* having the maximal *π*-rank among *k*-mers with the projection *s*^*k*^. Let *v* be a window with the projection 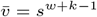 such that *v*[0, *k*) = *u* and *υ* [1, *k*+1) = *u*^*′*^≠ *u*. Then *f*_*γ*_ picks the position 0 while `rank` _*π*_(*u*) *>* `rank` _*π*_(*u*^*′*^). Hence *f* ≠ (*π, w*); the contradiction proves that *f*_*γ*_ is not a minimizer. □

#### S5.2. k-extension

Let *f* = (*ρ, w*) be a minimizer with the parameters (*σ, k, w*) and *k*^*′*^ *> k* be an integer. The local scheme *f*^*′*^ with the parameters (*σ, k*^*′*^, *w*) is defined as follows: in every (*w* + *k*^*′*^ − 1)-window *υ, f*^*′*^ picks the position chosen by *f* in the (*w* + *k* − 1)-prefix of *υ*.

##### Lemma 3

If *w > k*^*′*^, then *f*^*′*^ is not a minimizer.

*Proof* Suppose that *f*^*′*^ = (*ρ*^*′*^, *w*) is a minimizer. Let 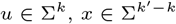 be such that `rank` _*ρ*_*′* (*ux*) = 0. As a minimizer, *f*^*′*^ picks the starting position of *ux* in every (*w* + *k*^*′*^ − 1)-window in which this *k*^*′*^-mer occurs. As *w > k*^*′*^ *> k*, every two *k*-mers appear in some (*w* + *k*^*′*^ − 1)-window; hence `rank` _*ρ*_(*u*) = 0.

Consider some *k*^*′*^-mer *uy* υ *ux*. Since *w > k*^*′*^, there exists a (*w* + *k*^*′*^ − 1)-window *v* with the prefix *uyux*. Since `rank` _*ρ*_(*u*) = 0, *f*^*′*^ by definition picks the position 0 of *υ*, while *υ* contains the *k*^*′*^-mer *ux* of smaller *ρ*^*′*^-rank. This contradiction proves that *f*^*′*^ is not a minimizer. □

If *w* ≤ *k*^*′*^, the answer to the question whether *f*^*′*^ is a minimizer depends on both *w* and *ρ*. In particular, if there exist (*w* +*k*^*′*^ −1)-windows *υ*_1_ and *υ*_2_ such that (i) *υ*_1_ begins with the *k*^*′*^-mer *ux* and contains the *k*^*′*^-mer *uy*; (ii) *υ*_2_ begins with *uy* and contains *ux*; (iii) *u* has the lowest *ρ*-rank among all *k*-mers in the (*w* + *k* − 1)-prefixes of *υ*_1_ and *υ*_2_, then *f*^*′*^ is not a minimizer. Indeed, similar to Lemma 3, *f*^*′*^ picks the position 0 in both *υ*_1_ and *υ*_2_, and thus *ux* and *uy* cannot be compared.

The authors of [de Blasio et al., Practical universal *k*-mer sets for minimizer schemes. ACM-BCB 2019] used the following notion of extension: if *ρ* and *ρ*^*′*^ are orders of *k*-mers and (*k* + 1)-mers respectively, and for every *u, u*^*′*^ ∈ Σ^*k*^, *a, b* ∈ Σ the inequality `rank` _*ρ*_(*u*) *<* `rank` _*ρ*_(*u*^*′*^) implies 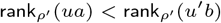, then *ρ*^*′*^ is an extension of *ρ*. They claimed, in Theorem 3 of the cited paper, that 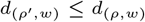 for any extension *ρ*^*′*^ of *ρ*. Their proof has a flaw: they claim that if the minimizer (*ρ*^*′*^, *w*) picks position *i* in some (*w* + *k*)-window *v* due to a (*k* + 1)-mer *ua* at this position, then the minimizer (*ρ, w*) picks the same position *i* in *υ* [0, *w* + *k*− 1) due to the *k*-mer *u*. But *υ* can contain *ub*, where *b* ≠ *a* and 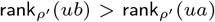, in a position *j < i*. In this case, (*ρ, w*) does not pick the position *i* in *υ* [0, *w* + *k* − 1), because ties are broken to the left.

The flawed proof does not imply automatically that Theorem 3 of [de Blasio et al.] fails, but our experiments show that the application of the particular extension defined in Section 3.3 in some cases increases density, so the theorem in the stated form is indeed incorrect.

### S6. Proof of Theorem 3

We call a window *v* ∈ (Σ *×* Γ)^*w*+*k*^ *bad* if it is charged by *ρ×τ* while *υ*_Σ_ is free for *ρ*, and *good* if *υ* is free for *ρ×τ* while *v*_Σ_ is charged by *ρ*. Let *B* and *G* denote the sets of bad and good windows, respectively. The order *ρ × τ* charges exactly *γ*^*w*+*k*^ *c* + |*B*| − |*G*| windows, where *c* is the number of windows charged by *ρ*. Consequently, 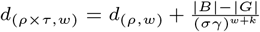. Nesxt, let *B*^←^ and G^?^ be the sets of bad and good windows for the order *ρ*^←^*× τ*, respectively.

#### Claim 1

|*B*| ≤ |*G*^←^ |.

*Proof of the Claim* Let *υ* ∈ *B* and let *u* be the minimum-rank *k*-mer of *υ*. Then, *u*_Σ_ has the minimum *ρ*-rank among the *k*-mers of *υ*_Σ_. By the definition of bad, *υ*_Σ_ is not charged by *ρ*, so *u*_Σ_ is neither a prefix nor a unique suffix of *υ*_Σ_. Hence, *u* is a suffix of *υ*, and *u*_Σ_ occurs more than once in *υ*_Σ_. Moreover, `rank` _*τ*_ (*u*_Γ_) *<* `rank` _*τ*_ (*z*_Γ_) for each other *k*-mer *z* of *υ* such that *z*_Σ_ = *u*_Σ_. We define the (*w*+*k*)-window *x* = *ϕ*(*υ*) as follows: 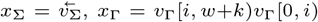, where *i* is the starting position of the leftmost occurrence of *u*_Σ_ in *υ*_Σ_ (Figure S1).

By the definition of mirror, the prefix 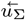 of *x*_Σ_ has the minimum *ρ*^←^-rank among the *k*-mers of *x*_Σ_. Then, *x*_Σ_ is charged by *ρ*^←^. Let *z* = *υ* [*i, i* + *k*), *u*^*′*^ = *x*[0, *k*), *z*^*′*^ = *x*[*w*−*i, w*−*i*+*k*), and *y* = *x*[*w, w* + *k*). Since *z*_Σ_ = *u*_Σ_ by the choice of *i*, we have `rank` _*τ*_ (*u*_Γ_) *<* `rank` _*τ*_ (*z*_Γ_). Since 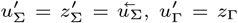 and 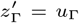, we get 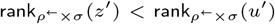. Note that 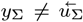 because *u*_Σ_ is not a prefix of *υ*_Σ_. As 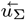 has the minimum *ρ*^←^-rank among the *k*-mers of *x*_Σ_, we have 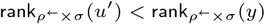. Since *z*^*′*^ has smaller rank than both the *k*-prefix and the *k*-suffix of *x, x* is free for *ρ*^←^ *× τ*. Then, *x* ∈ *G*^←^ by the definition of a good window. Hence, *ϕ* maps *B* to *G*^←^.

**Fig. S1.**
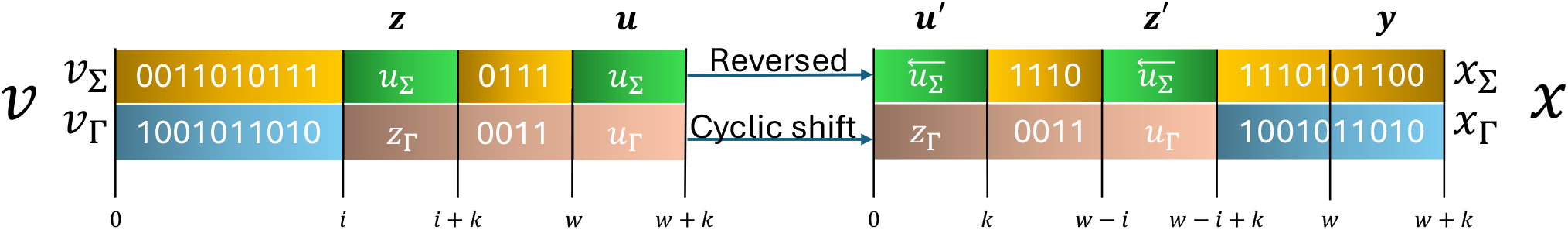
Example of mapping a bad (for *ρ × τ*) window *v* to a good (for *ρ*^←^ *× τ*) window *x*.

Now let *υ, υ*^*′*^ ∈ *B, υ* ≠*υ*^*′*^, *x* = *ϕ*(*υ*), *x*^*′*^ = *ϕ*(*υ*^*′*^). If 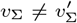, then 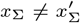. If 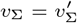, then 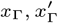 are obtained by the same shift of *υ*_Γ_ and 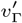, respectively. Since *υ* ≠*υ*^*′*^, we have 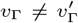, and then 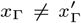. Therefore, in both cases *x* ≠ *x*^*′*^. Thus, we proved that *ϕ* is injective. The claim now follows. □

Since (*ρ*^←^)^←^ = *ρ*, the claim implies |*B*^←^ | ≤ |*G*|. Then

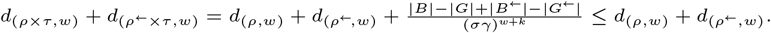

The result now follows by Lemma 1.

### S7. Proof of Theorem 4

Similar to the proof of Theorem 3, we define bad and good windows. For *υ* ∈ Σ^*w*+*k*^ and *a* ∈ Σ, the window *va* is bad (good) for *ρ*_1_ if it is charged (not charged) by *ρ*_1_ while *υ* is not charged (charged) by *ρ*. By *B*_1_ and *G*_1_ we denote the sets of bad and good windows for *ρ*_1_, respectively. Similarly, we define bad and good windows *av* for *ρ*_2_ and denote their sets by *B*_2_ and *G*_2_, respectively. We prove the analog of the claim from Theorem 3.

#### Claim 2

|*B*_1_| ≤ |*G*_2_| and |*B*_2_| ≤ |*G*_1_|.

*Proof of the Claim* Let *va* ∈ *B*_1_ and let *ub* be the (*k*+1)-mer in *υa* of minimum *ρ*_1_-rank (here *a, b* ∈ Σ). By the definition of *ρ*_1_, *u* has the minimum *ρ*-rank in *υ*. Since *υ* is free for *ρ, u* is not a prefix of *υ*. Hence, *ub* is a unique suffix of *υa*; in particular, *b* = *a*. Then, *u* is a suffix of *υ* and has other occurrences in *υ* because *υ* is free for *ρ*. Thus, *v* contains a (*k*+1)-mer *ua*^*′*^, where *a*^*′*^ *> a* as `rank` _*ρ*_ (*ua*^*′*^) *>* `rank` _*ρ*_ (*ua*).

The window 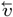 is charged by *ρ*^←^ as its minimum-rank *k*-mer 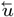 is its prefix (but not suffix). Consider the window 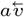. Its prefix 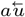 has smaller *ρ*_2_-rank than its (*k*+1)-suffix, as this suffix does not end with 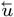. Further, 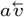 contains the (*k*+1)-mer 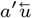 with *a*^*′*^ *> a* and thus 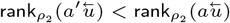). Hence, 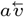 is free for *ρ*_2_. We thus proved 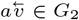. As the reversal of every window from *B*_1_ belongs to *G*_2_, we have |*B*_1_| ≤ |*G*_2_|. A symmetric argument proves |*B*_2_| ≤ |*G*_1_|, completing the claim. □

Let *c, c*^←^ be the numbers of windows charged by *ρ* and *ρ*^←^, respectively. Then, *ρ*_1_ charges *σc* + |*B*_1_| − |*G*_1_| windows, and *ρ*_2_ charges *σc*^←^ + |*B*_2_|−|*G*_2_| windows. Hence, 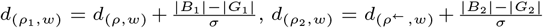. Substituting the inequalities from the claim, we get

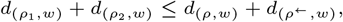

implying the theorem in view of Lemma 1.

### S8. GreedyMini workflows

**Fig. S2.**
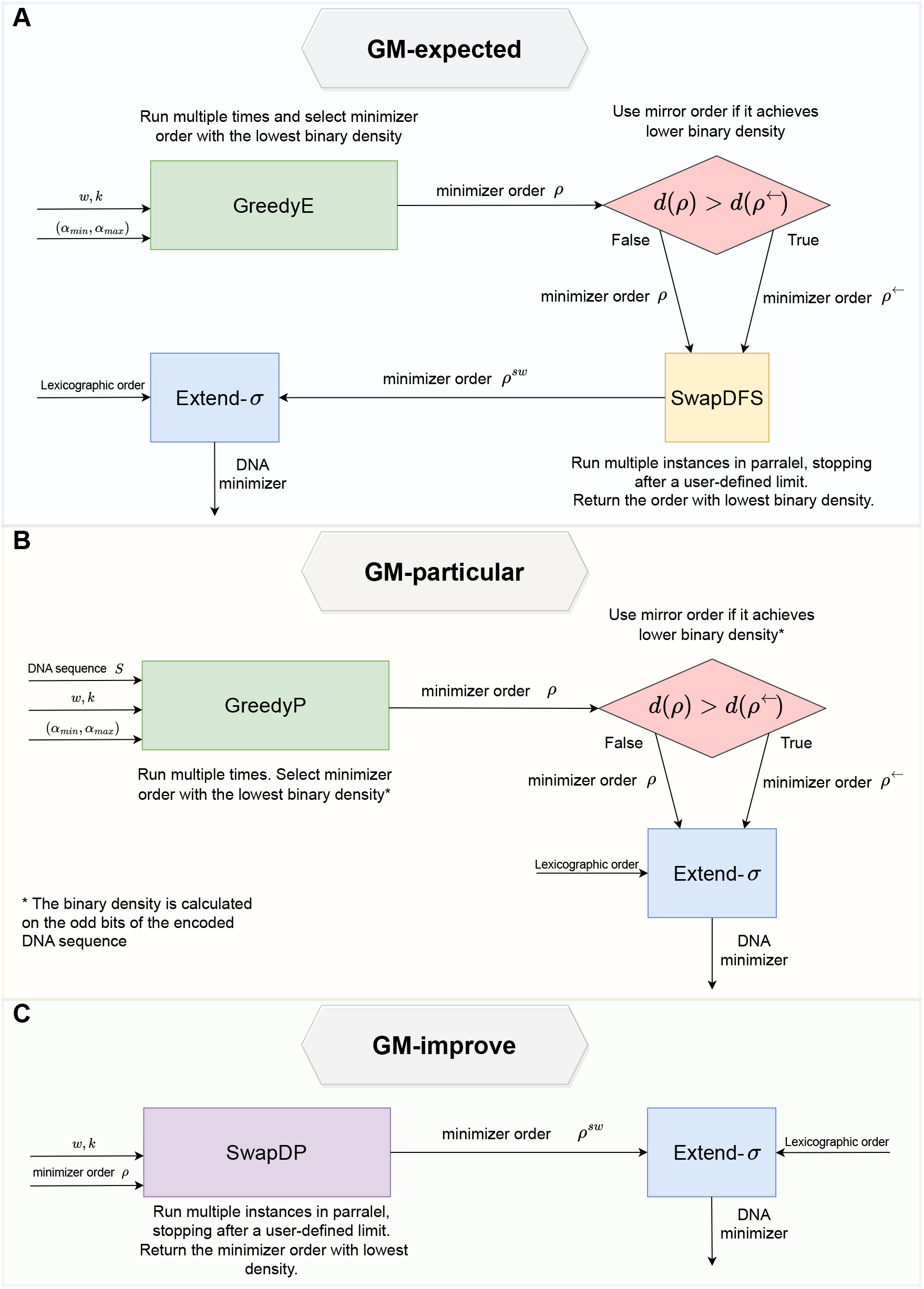
Workflow illustration of (A) GM-expected pipeline to generate a low-expected-density DNA minimizer, (B) GM-particular pipeline to generate a low-particular-density DNA minimizer, and (C) GM-improve pipeline to improve a (binary) minimizer to larger *w* values.

### S9. Running DeepMinimizer and polar sets

We failed to run DeepMinimizer [11] and polar sets [22] due to errors received when running their code. Both implementations contained unspecified file hierarchies, hard-coded paths, and lacked essential documentation, such as the required dependencies and their versions. For instances where hard-coded paths were present, we replaced them with local paths. After adjusting the paths, the following issues arose:

#### Polar sets

We ran the following command:

~~~
python anchor_distributed.py sequence_1M -w 12 --kmin 3
--kmax 16 -p 10
~~~

It raised the following error:

~~~
Traceback (most recent call last):
  File “/home/dsi/tzionyi/polarset/anchor_distributed.py”,
  line 18, in <module> assert working_dir is not None
~~~

#### DeepMinimizer

We ran the following command:

~~~
python run_scripts.py
~~~

It raised the following error:

~~~
RuntimeError: one_hot is only applicable to index tensor.
~~~

### S10. Construction runtime and maximum memory usage of GM-expected and GM-particular

We ran `GM` -`expected` and `GM` -`particular` on a Linux server equipped with 2 *×* Intel(R) Xeon(R) Gold 6338 CPUs (total cores 64) @ 2.00GHz and 512GB of RAM. We utilized 64 available cores. We report the construction runtime (Supplementary Figure S3A) and maximum memory usage (Supplementary Figure S3B) of our implementation of `GM` -`expected` over various combinations of *w* and *k* without extending *k* (Theorem 4).

The measured construction runtimes fit the bound of `GreedyE` given by Theorem 1 (*O*(*σ*^2*k*^ + *wσ*^*k*+*w*^)). Focusing on *w* + *k >* 18 to avoid effects of small values, we observe for *k >* 2*w* an exponential dependence on *k*, while for *k* ≤ 2*w* we observe an exponential dependence on *w* + *k*. Note that we run the local search by `Swap` iterations with a time limit according to the runtime of `GreedyMini`, and hence their runtimes are equal. The maximum memory usage matches that of `GreedyE` and `Swap` by Theorems 1 and 2 (*O*(*σ*^*k*^ + *w*)). Note that due to the utilization of various libraries and other computational factors, our implementation of `GM` -`expected` has a baseline memory usage, and thus the increase in maximum memory usage is only observed for *k* ≥ 14.

In addition, we report the construction runtime (Supplementary Figure S4A) and maximum memory usage (Supplementary Figure S4B) of our implementation of `GM`-`particular`. Over a sequence length of 1 million nucleotides, the construction runtime increased exponentially with *k* for *k >* 12, and increased much more slightly with *w*. The maximum memory usage increased with *w* more profoundly than with *k*.

**Fig. S3.**
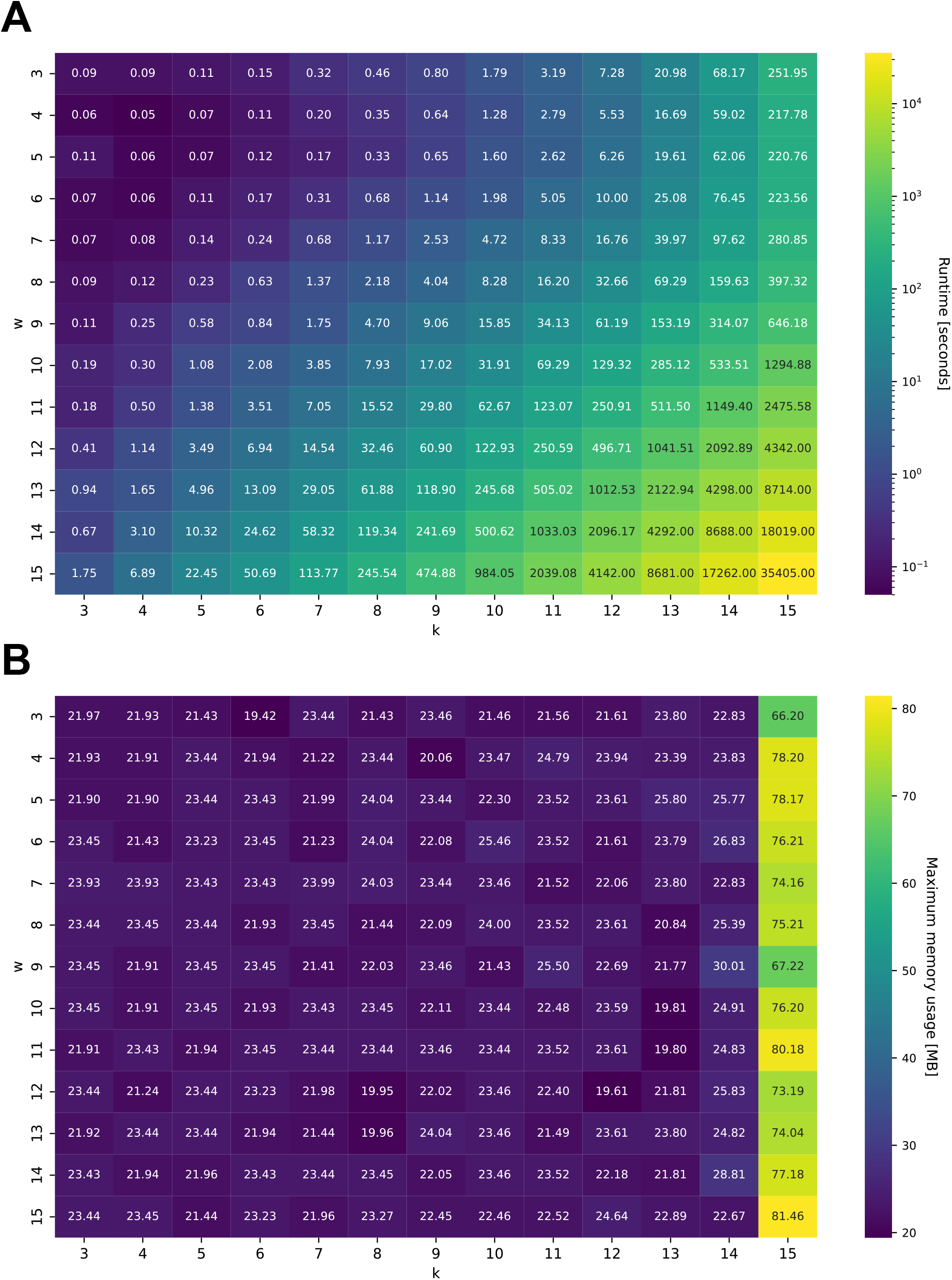
Runtime (A) and maximum memory usage (B) of GM-expected on various *w* and *k* values.

**Fig. S4.**
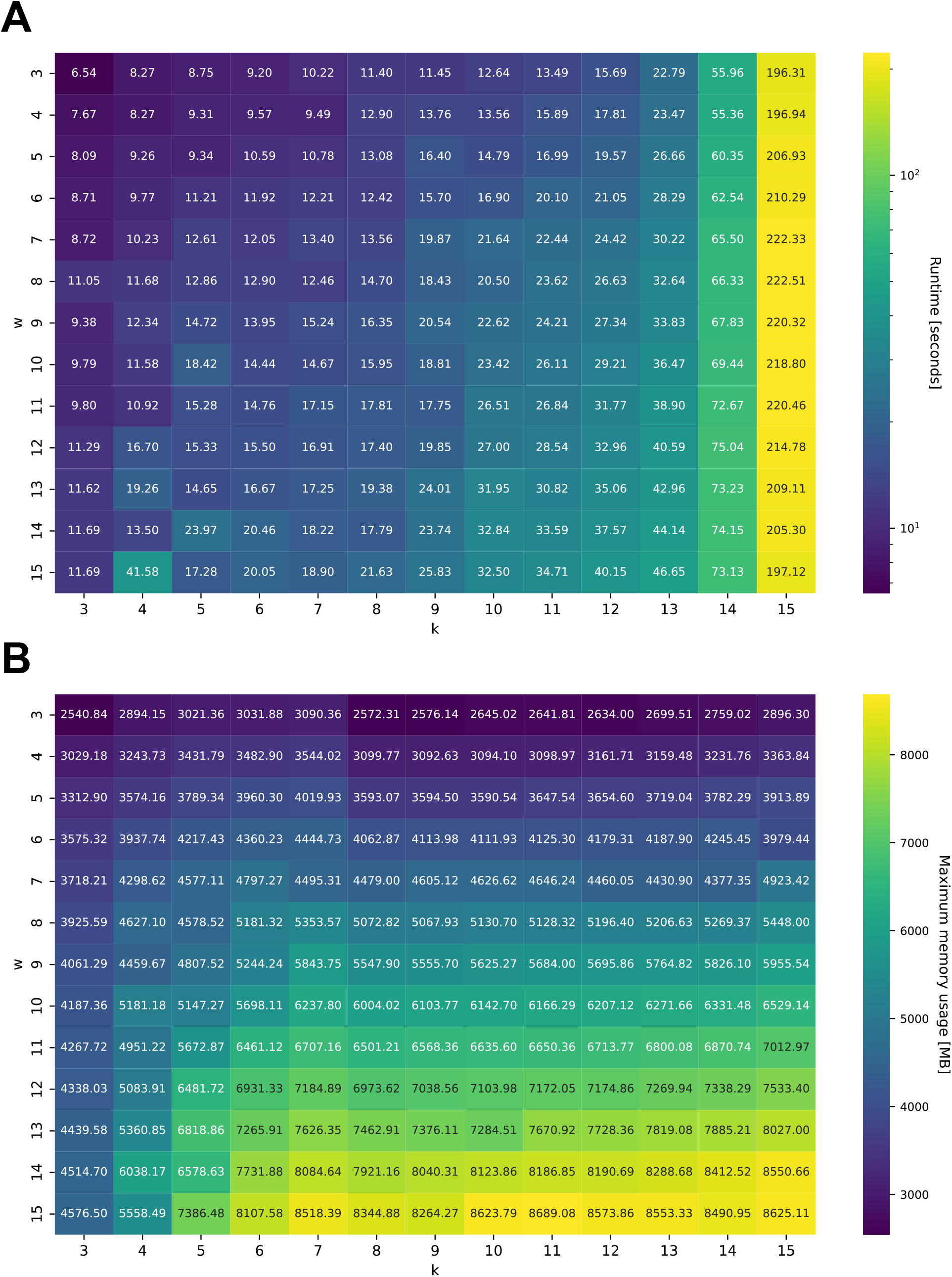
Runtime (A) and maximum memory usage (B) of GM-particular on various *w* and *k* values over 1 million nucleotides from Chromosome X.

